# Heterogeneity and molecular programming of progenitors for motor neurons and oligodendrocytes

**DOI:** 10.1101/2021.06.01.446521

**Authors:** Lingyan Xing, Rui Chai, Jiaqi Wang, Jiaqi Lin, Hanyang Li, Yueqi Wang, Biqin Lai, Junjie Sun, Gang Chen

**Affiliations:** Key Laboratory of Neuroregeneration of Jiangsu and the Ministry of Education, Co-innovation Center of Neuroregeneration, NMPA Key Laboratory for Research and Evaluation of Tissue Engineering Technology Products, Nantong University, Nantong, China; Department of Physiology, School of Medicine, Nantong University, Nantong, China; School of Medicine, University of Utah, Salt Lake City, USA; Key Laboratory for Stem Cells and Tissue Engineering (Sun Yat-sen University), Ministry of Education, Co-innovation Center of Neuroregeneration Nantong University Nantong 226001 China; Basic Medical Research Center, School of Medicine, Nantong University, Nantong, China

**Keywords:** single-cell RNA sequencing (scRNA-seq), zebrafish, heterogeneity, oligodendrocyte progenitor cells (OPCs), neural progenitors, motor neurons, pMN domain, spinal cord

## Abstract

The pMN domain is a restricted domain in the ventral spinal cords, defined by the expression of *olig2* gene. The fate determination of pMN progenitors is highly temporally and spatially regulated, with motor neurons and oligodendrocyte progenitor cells (OPCs) developing sequentially. Insight into the heterogeneity and molecular programs of pMN progenitors is currently lacking. With the zebrafish model, we identified multiple states of neural progenitors using single-cell sequencing: proliferating progenitors, common progenitors for both motor neurons and OPCs, and restricted precursors <u>for</u> either motor neurons or OPCs. We found specific molecular programs for neural progenitor fate transition, and manipulations of representative genes in the motor neuron or OPC lineage confirmed their critical role in cell fate determination. The transcription factor NPAS3 is necessary for the development of the OPC lineage and can interact with many known genes associated with schizophrenia. Deciphering progenitor heterogeneity and molecular mechanisms for these transitions will elucidate the formation of complex neuron-glia networks in the central nervous system during development, and understand the basis of neurodevelopmental disorders.

## Introduction

In development, neural progenitors have binary fate choices. They sequentially generate neurons and glia, which is the basis of complexity in the central nervous system (CNS). In the spinal cord, both motor neurons and oligodendrocytes arise from the pMN domain; progenitors in this domain are defined by the gene marker *olig2*, and sequentially form motor neurons and oligodendrocyte progenitor cells (OPCs). This process is both temporally and context regulated. Neurogenin1 and Neurogenin2 are upregulated preceding neurogenesis but downregulated in oligodendrogenesis (Mizuguchi, Sugimori, Takebayashi, Kosako, Nagao, Yoshida, Nabeshima, Shimamura, and Nakafuku 2001). OPC specification is initiated with the induction of *sox10*, which leads to the upregulation of *nkx2.2* (Zhou, Choi, and Anderson 2001).

Though multiple studies have revealed several key factors in the development of motor neurons and OPCs, the transcriptional signatures of progenitors and their daughter cells are not fully understood. In addition, the heterogeneity of motor neuron and OPC progenitors is controversial (Ravanelli and Appel 2015). For example, individual progenitors grown in culture or traced by recombinant retroviral infection or fluorescent dye can generate both neurons and glia in flies, mice, or zebrafish (Schmid, Chiba, and Doe 1999; Qian, Shen, Goderie, He, Capela, Davis, and Temple 2000; Leber and Sanes 1995; Park, Shin, and Appel 2004). However, clonal lineage testing and live imaging reveal that subsets of progenitors are fate-restricted, forming either motor neurons or glial cells in mice or zebrafish (Qian, Goderie, Shen, Stern, and Temple 1998; Ravanelli and Appel 2015). These findings from tissue transplantation, lineage tracing, or cell ablation have been heavily based on reporter genes, the non-specificity of which may lead to confounding results. The single-cell RNA sequencing (scRNA-seq) approach offers a high resolution method to decipher the molecular programs driving the transition of neural fates. Recent studies have used large-scale profiling of single cells in zebrafish (Wagner, Weinreb, Collins, Briggs, Megason, and Klein 2018; Farrell, Wang, Riesenfeld, Shekhar, Regev, and Schier 2018), but these studies focus on early embryogenesis, before most glial cells are specified.

Here we applied the scRNA-seq technique to sequence 11920 cells in zebrafish ventral spinal cords, identify temporal lineages, and decode the transcriptome landscapes for progenitor fate transitions. Our study identified heterogeneity in the molecular signatures of pMN progenitors, and reveals that both motor neurons and OPCs derive from common pMN progenitors but with distinct lineage-restricted precursors. We also demonstrate that *myt1* (Myelin Transcription Factor 1) is necessary for motor neuron specification, and that *npas3* (Neuronal PAS Domain Protein 3) is necessary for development of the OPC lineage. Interestingly, with two published datasets, we confirm that *npas3* is restricted to the OPC lineage. As a transcription factor, NPAS3 may have interaction with known schizophrenia-associated genes. These provide new insights into the role of *npas3* in neural circuit formation and neuropsychiatric disorders. Our study unveils novel molecular programming for neuron/glia differentiation and suggests approaches for multiple applications, including directed cell differentiation or reprogramming. Notably, we also established open web-based datasets for researchers to access.

## Results

### Single-cell sequencing identifies distinct cell populations in the pMN domain

Progenitors in the ventral spinal cords (pMN domains) of zebrafish initiate differentiation into motor neurons prior to 24 hours post fertilization (hpf), and motor neuron generation is largely accomplished by 48-51 hpf (Reimer, Norris, Ohnmacht, Patani, Zhong, Dias, Kuscha, Scott, Chen, Rozov, Frazer, Wyatt, Higashijima, Patton, Panula, Chandran, Becker, and Becker 2013). At 36 hpf, sox10 and nkx2.2 positive cells (OPC lineage cells) have been observed (Mathews and Appel 2016). To identify the transcriptome profiles in pMN progenitors and their offspring, we applied fluorescence activated cell sorting (FACS) to specifically sort cells labeled by the transgenic line *Tg (olig2:dsRed)* (dsRed is expressed under the control of the olig2 regulatory element) (Shin, Park, Topczewska, Mawdsley, and Appel 2003) at 42 hpf, a stage when pMN progenitors, motor neurons, and OPC-related cells are all labeled with dsRed (Supplemental Figure 1A-D). To specifically identify progenitors and their offspring in the ventral spinal cords, we only retained the fish trunks for single cell isolation (Figure 1A). ∼200 fish trunks were collected to minimize individual variation. Altogether, we transcriptionally profiled 11920 cells with the high-throughput single-cell systems: 10X Genomics (Zheng, Terry, Belgrader, Ryvkin, Bent, Wilson, Ziraldo, Wheeler, McDermott, Zhu, Gregory, Shuga, Montesclaros, Underwood, Masquelier, Nishimura, Schnall-Levin, Wyatt, Hindson, Bharadwaj, Wong, Ness, Beppu, Deeg, McFarland, Loeb, Valente, Ericson, Stevens, Radich, Mikkelsen, Hindson, and Bielas 2017). A median of 27796 reads per cell were detected, corresponding to a median of 1261 genes per cell (Supplemental Figure 2A-C). This depth is comparable to similar studies (Zheng, Terry, Belgrader, Ryvkin, Bent, Wilson, Ziraldo, Wheeler, McDermott, Zhu, Gregory, Shuga, Montesclaros, Underwood, Masquelier, Nishimura, Schnall-Levin, Wyatt, Hindson, Bharadwaj, Wong, Ness, Beppu, Deeg, McFarland, Loeb, Valente, Ericson, Stevens, Radich, Mikkelsen, Hindson, and Bielas 2017).

**Figure 1.**
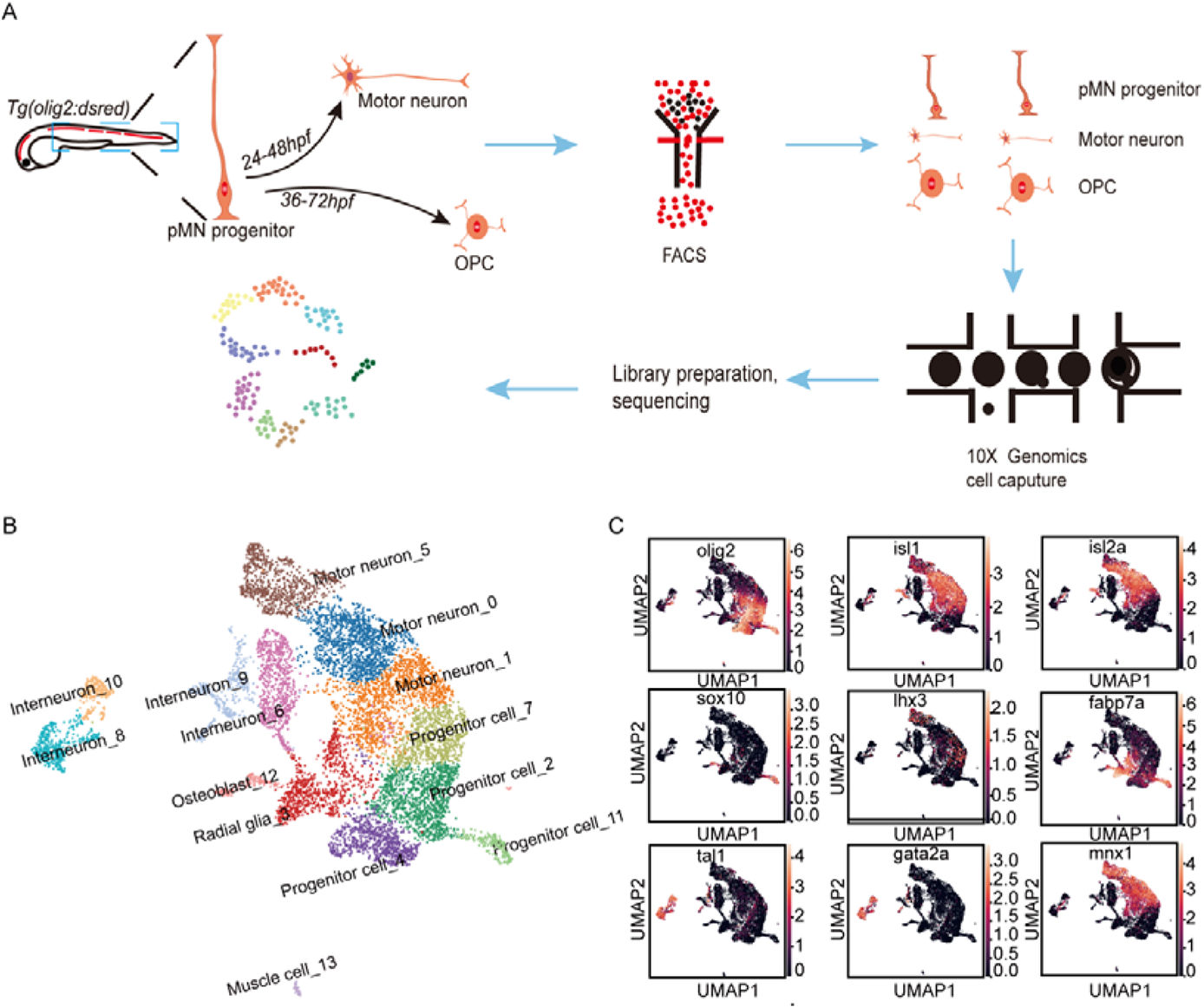
Clustering of sorted cells with scRNA-seq. A) Diagrams of the scRNA-seq workflow. B) UMAP clustering of single-cell samples. Colors indicate distinct groups. C) The expression of marker genes visualized by UMAP. The color denotes the range of log2 transformed fragments per kilobase of transcript per million reads mapped (FPKM) expression values. The darker the color, the lower the expression.

Sorted cells were classified based on their gene expression pattern. RNA-seq expression patterns were projected into UMAP (Uniform Manifold Approximation and Projection) or t-SNE (t-distributed Stochastic Neighbor Embedding), a nonlinear dimensionality reduction algorithm, for cell population annotation (Becht, McInnes, Healy, Dutertre, Kwok, Ng, Ginhoux, and Newell 2018). All together, 13 distinct cell clusters were identified (Figure 1B, Supplemental Figure 2D).

By combining known gene markers in progenitors, neurons, or glia in the spinal cords, we defined these cells: pMN progenitors, motor neurons, radial glia, and interneurons. pMN progenitors were defined by their expression of *olig2* (Park, Shin, and Appel 2004). Motor neurons were enriched for *mnx2a*, *mnx2b*, *mnx1*, *lhx3*, or *isl1* (Seredick, Ryswyk, Hutchinson, and Eisen 2012). Radial glia were defined by fabp7 and GFAP (Lange, Rost, Machate, Reinhardt, Lesche, Weber, Kuscha, Dahl, Rulands, and Brand 2020). Finally, V2b interneurons were marked by *gata3*, *tal1*, and *gata2a*, V2a interneurons by *sox14* and *vsx2*, V3 interneurons by *uncx*, and V0 interneurons by *evx1-1* and *evx2* (Andrzejczuk, Banerjee, England, Voufo, Kamara, and Lewis 2018; Juarez-Morales, Schulte, Pezoa, Vallejo, Hilinski, England, de Jager, and Lewis 2016; Clovis, Seo, Kwon, Rhee, Yeo, Lee, Lee, and Lee 2016) (Figure 1B-C, Supplemental Figure 3A). These results were in line with the cell types labeled by this transgenic line in previous work (Takada and Appel 2010), confirming the reliability of our sequencing data. We also integrated the specific markers to identify cell clusters sorted by FACS but not mentioned previously. For example, the *tnnt3* and *tnnc2* genes marked muscle cells, and *meox1* and *postnb* identified osteoblasts (Yang, Shih, and Xu 2014; Filtz, Emery, Lu, Forster, Karasch, and Hallstrom 2015) (Supplemental Figure 3B).

### Diversity of pMN progenitors or precursors

To probe the relationships between the cells isolated by FACS, we performed partition-based graph abstraction (PAGA) analysis (Wolf, Angerer, and Theis 2018), which showed the topography and lineage of the cell clusters (Figure 2A-B). Interestingly, a few cell clusters, e.g. clusters 12 and 13, were disconnected from the *olig2*^+^ cells (Figure 2C), indicating that these cells were either from non-specific labeling of the *olig2* reporter or non-specific sorting by FACS. Therefore, we only retained the cells having a strong association with *olig2* progenitors for further analysis (Figure 2C, D).

**Figure 2.**
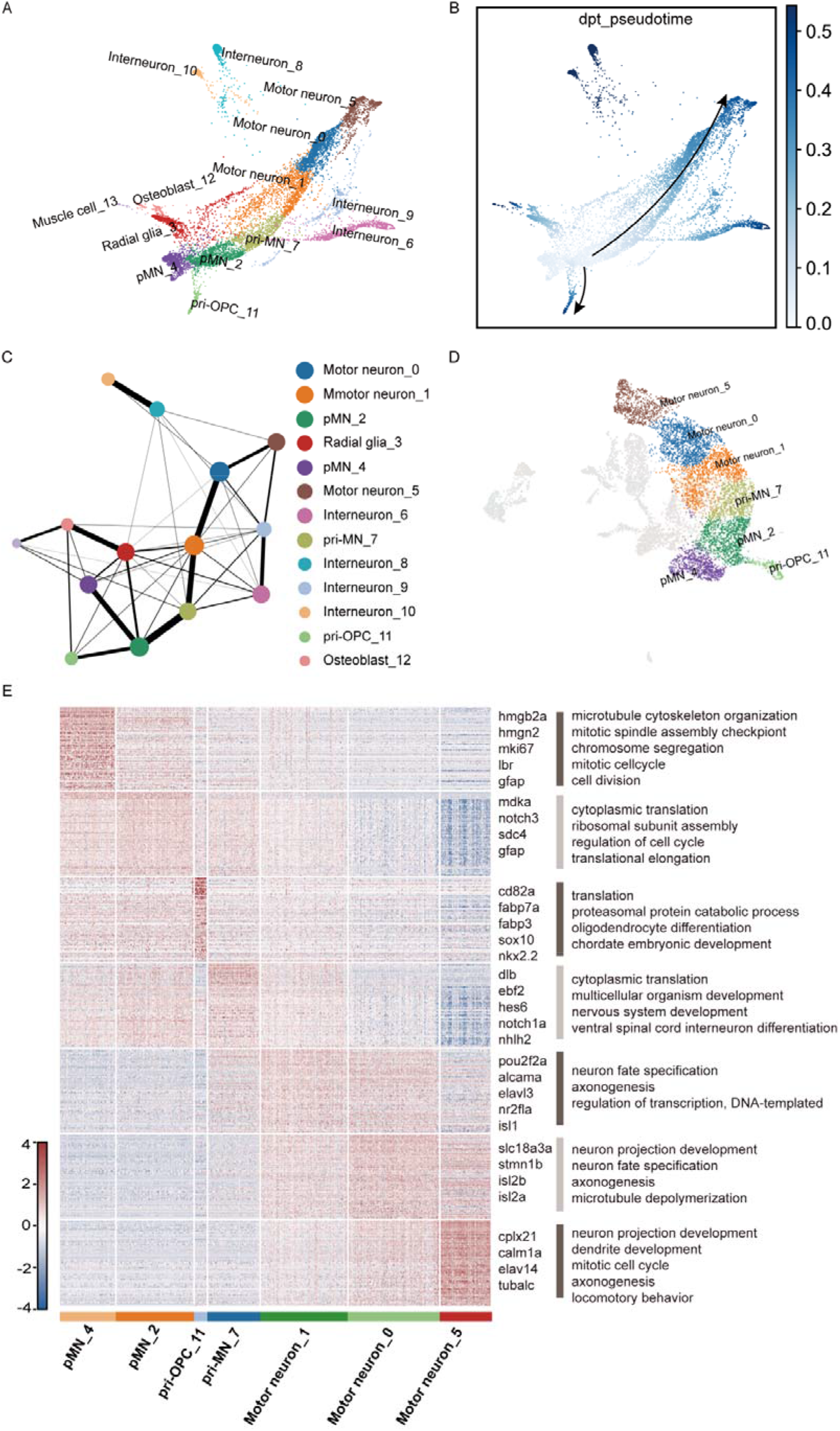
Cell clustering analyses of sorted cells. A) Cell trajectory analysis by Partition-based graph abstraction (PAGA). B) Pseudotemporal ordering of cells by PAGA. The darker color depicts more mature cells, and the lighter color denotes younger cells. C) Connections between cell types indicated by PAGA. Nodes denote subpopulations of cells, and edge weights depict the confidence of the connections between each set of two clusters. D) UMAP visualization of selected cells with strong connections. E) Heatmaps of signature genes and enriched GO terms in each subcluster.

These retained clusters encompass three types of motor neurons and four types of progenitors (*olig2*^+^). In the four subpopulations of pMN progenitors, precursors 7 and 11 branched into two trajectories (Figure 2A). Cluster 7 is highly enriched with the neuronal markers *nhlh2*, *mnx1*, and *isl1*, as well as the proneuronal markers *hes6, btg2,* and *ascl1* (Figure 1C, Figure 2E, Figure 3A-C), indicating that these cells were primitive motor neurons (pri-MNs). Consistently, this subpopulation transitioned into motor neurons (Figure 2A). In these different subpopulations of motor neurons (motor neuron_1, motor neuron_0, and motor neuron_5), the expression levels of known neuronal markers varied. For example, early motor neuronal marker *isl1* was expressed in motor neuron_1, *isl2b*, *isl2a*, and *elavl4* were enriched in motor neuron_0 and motor neuron_5, and mature neuronal markers *tuba1c* and *synap25a* were highly expressed in motor neuron_5. Consistently, the enriched pathways of highly expressed genes switched from neuronal specification to dendritic, axongenesis, and synaptogenesis in motor neuron_1 and motor neuron_5, respectively (Figure 2E). Consistent with our lineage paradigm, these cells were motor neurons at different developmental stages, from immature neurons to mature neurons (Figure 2A, E). Interestingly, we also found several novel markers expressed in subsets of these motor neurons, for example immature motor neurons (motor neuron_0) were enriched for *slc18a3a* and *stmn1b*, and mature neurons (motor neuron_5) were enriched for *cplx21* and *calm1a*. Notably, zebrafish show expression of genes which do not have mammalian orthologs; for example, *dlb* (delta-like protein B) was enriched in zebrafish immature neurons (motor neuron_1) (Figure 2E). Therefore, sc-RNAseq revealed that motor neurons underwent a complex developmental transition from the intermediate precursors (pri-MN) into mature neurons.

**Figure 3.**
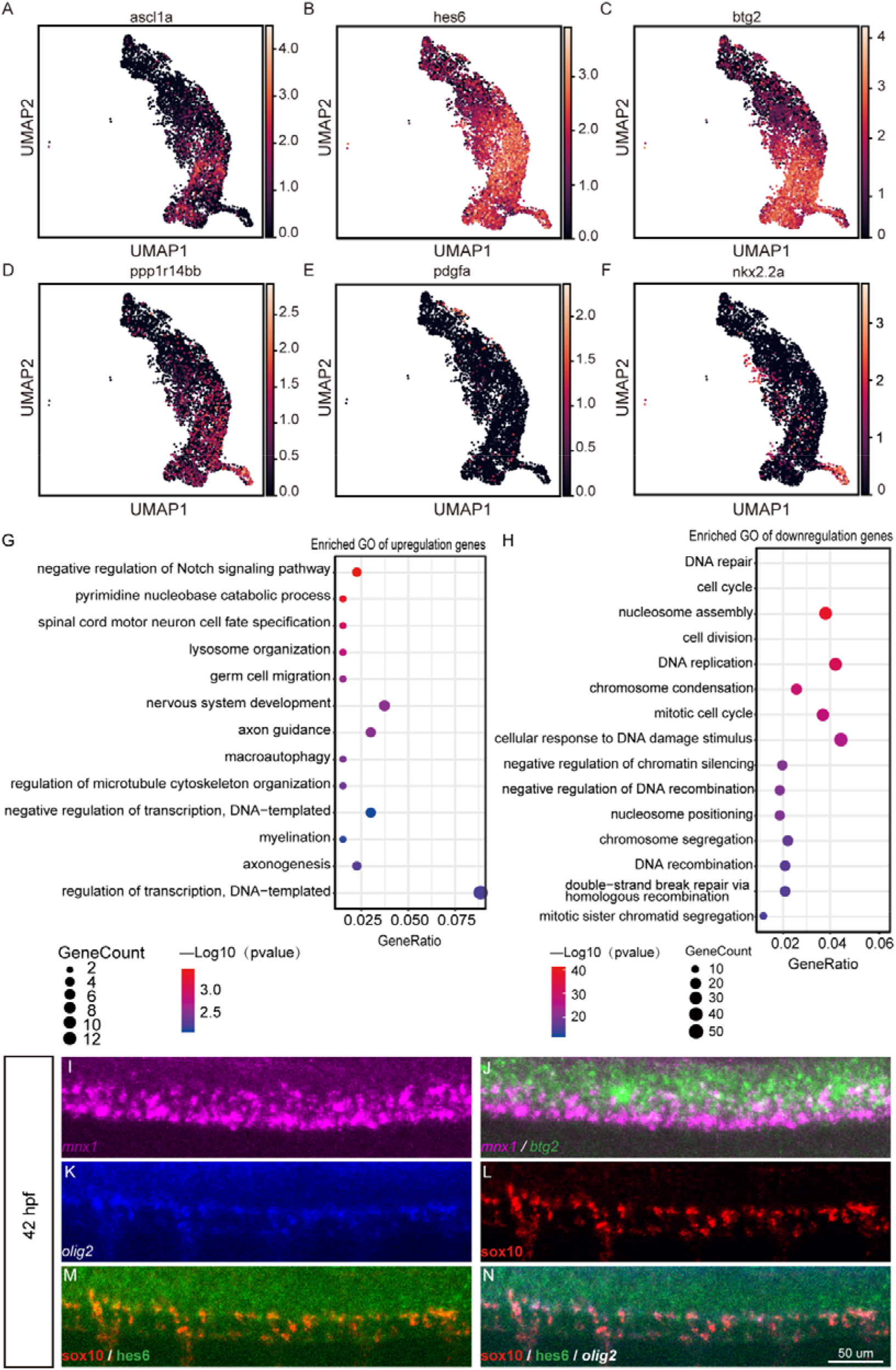
Identification of pMN progenitors and precursors. A-D) UMAP of key genes in progenitor or precursor cells. E-F) UMAP of key genes in OPC-lineage cells. G-H) GO terms enriched in pMN progenitors. Bubble sizes represent fold enrichment; color scales denote the statistical significance. Fisher’s exact test and the Bonferroni correction were performed to calculate p-values which were log transformed. I-J) *in situ* fluorescent hybridization of precursor markers *btg2* and motor neuron marker *mnx1* reveals the existence of pri-MNs in whole-mount embryos at 42 hpf. K-N) *in situ* fluorescent hybridization of precursor markers *hes6* and OPC marker *sox10/olig2* reveals the existence of pri-OPCs in whole-mount embryos at 42 hpf.

The precursor_11 in the other trajectory was enriched for the OPC markers *sox10* and *nkx2.2*, as well as progenitor or precursor markers including *hes6*, *btg2*, and *ppp1r14bb* (Figure 3B-D, F), indicating that this subpopulation was comprised of primitive OPCs (pri-OPCs) rather than OPCs. Consistently, it had low expression levels of the OPC marker *pdgf*α (Figure 3E).

Fluorescent *in situ* hybridyzation (FISH) was applied to elucidate the intermediate states of MNs and OPCs. At 42 hpf, when motor neurons were actively generated (Ohnmacht, Yang, Maurer, Barreiro-Iglesias, Tsarouchas, Wehner, Sieger, Becker, and Becker 2016), many cells co-expressed the proneuronal markers *btg2* and the neuronal marker *mnx1* (Figure 3I-J), indicating that they were pri-MNs. Likewise, at 42hpf, we observed that almost all *sox10*+ cells or *sox10+olig2+* cells were co-expressed with *hes6*, indicating that these were pri-OPCs (Figure 3K-N). Cells we analyzed at 42 hpf, a stage when motor neurons are still actively generated, include pri-OPCs as well as pri-MN cells. The existence of pri-OPCs at this early developmental stage indicates that the motor neuron and the OPC lineages segregate early, and does not support the progenitors developing first into motor neurons and later switching to the OPC lineage.

The pMN_2 and pMN_4 clusters were enriched for *olig2*, and developed into two lineages, indicating these were pMN progenitor cells. From lineage analysis, both motor neuron and OPC restricted precursors appeared to be derived from the cluster pMN_2 rather than the pMN_4 cluster (Figure 2A). pMN_2 and pMN_4 progenitors had genes in common, such as GFAP, a neural progenitor marker (Figure 2E, Supplemental Figure 5A-B), as shown by live imaging of *Tg(olig2:dsRed; gfap:GFP)* animals, in which dsRed driven by the *olig2* promoter was co-expressed with *GFP* driven by the *gfap* promoter.

By differentially expressed gene (DEG) analysis, we found that multiple known proliferative markers were downregulated in pMN_2 compared to pMN_4, including *mki67*, *hmgb2a,* and *hmgb2b* (Supplemental Figure 4D-E). Notably, the transition from pMN_4 to pMN_2 was accompanied by dramatic down-regulation of expression of most genes, and upregulation in fewer genes (Supplemental Figure 4C). Go analysis shows that genes downregulated in pMN_2 were enriched in DNA repair, cell cycle, and chromosome-remodeling-related pathways (e.g. nucleosome assembly, chromosome condensation), indicating a decreased proliferative capacity in pMN_2. Go analysis for upregulated pathways in pMN_2 vs. pMN_4 revealed that transcriptional regulation and nervous system development pathways were enriched (Figure 3G-H).

Taken together, all of these results show that the pMN_4 cluster represents proliferating states while the pMN_2 cluster corresponds with differentiating states of neural progenitors; the transition between these two states was accompanied by dynamic transcriptional profiling changes, especially with the down-regulation of genes involved in the cell cycle or proliferation. Our results indicate that proliferation of progenitors counteracts differentiation; when the cell proliferation capacity is reduced, differentiation occurs. The number of downregulated DEGs in the differentiation state, which were enriched in chromatin remodeling and proliferation, consistently far exceeded the number of upregulated DEGs in our progenitor DEG analysis (Figure 3G-H, Supplemental Figure 4C).

### Lineage trajectories of pMN progenitors

The existence of pri-MNs and pri-OPCs indicates motor neurons and OPCs derive from distinct restricted precursors. However, whether pri-MNs and pri-OPCs originate from common olig2+ progenitors is unclear (Figure 4A-B). We observed that when PAGA lineage analysis was applied, both the motor neuron and OPC lineages were derived from the same cluster of pMN progenitors (Figure 2A). This was in line with the subclustering analysis of pMN progenitors and precursors (Supplemental Figure 5A-C). To confirm this conclusion, we used Monocle, another powerful algorithm based on transcriptional profiling (Trapnell, Cacchiarelli, Grimsby, Pokharel, Li, Morse, Lennon, Livak, Mikkelsen, and Rinn 2014), for cell lineage analysis. This algorithm gave us the same results, in which only the population of pMN progenitor 2 cells branched into both the motor neuron lineage (pri-MN 7) and the OPC lineage (pri-OPC 11) (Figure 4C-D).

**Figure 4.**
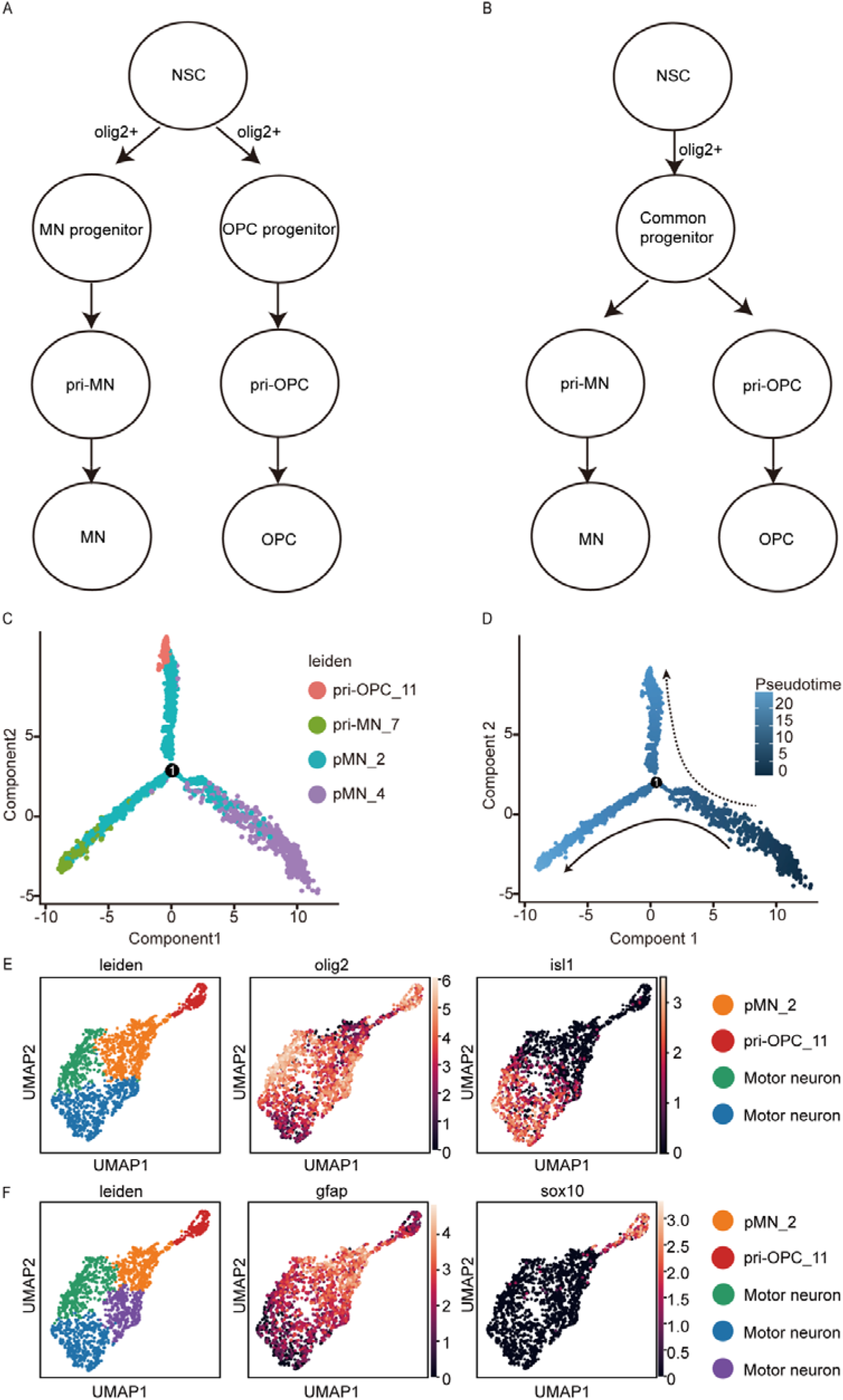
Lineage analyses of pMN progenitors and precursors. A-B) Possible models for the lineages of pri-MNs and pri-OPCs. C-D) Additional analysis of precursor fate determination into neurons and OPCs. Selected cells are ordered by pseudotime analysis (Monocle). Each dot represents an individual cell. E-F) Subclustering of pMN_2 progenitors and their offspring into four or five groups. Notice that the pMN_2 progenitors retain one cluster when subclustering into more groups.

To exclude the possibility that the pri-OPCs and pri-MNs are derived from distinct subpopulations of pMN_2, we subclustered pMN_2, pri-OPC, and pri-MN cells into four or five groups (Figure 4E-F) in an unbiased way, and found that the pMN_2 cells maintained a single cluster. This confirmed that the pri-OPC and the pri-MN cells originated from the same progenitors.

### Molecular programming for the MN and OPC lineages

So far, we have characterized four subpopulations of pMN progenitors or precursors. pri-MNs and pri-OPCs segregate from the pMN progenitors. Therefore, understanding the molecular programs that underlie the transition from pMN progenitors to pri-OPC or pri-MN will elucidate the cell fate decisions that ultimately determine motor neuron or OPC lineages. With this in mind, we utilized Monocle to analyze temporal changes in gene expression for MN and OPC fate determination (Trapnell, Cacchiarelli, Grimsby, Pokharel, Li, Morse, Lennon, Livak, Mikkelsen, and Rinn 2014). We selected these TFs among the top 500 differentially expressed genes (DEGs) that biased the fates of pMN_2 progenitors (Figure 5B).

**Figure 5.**
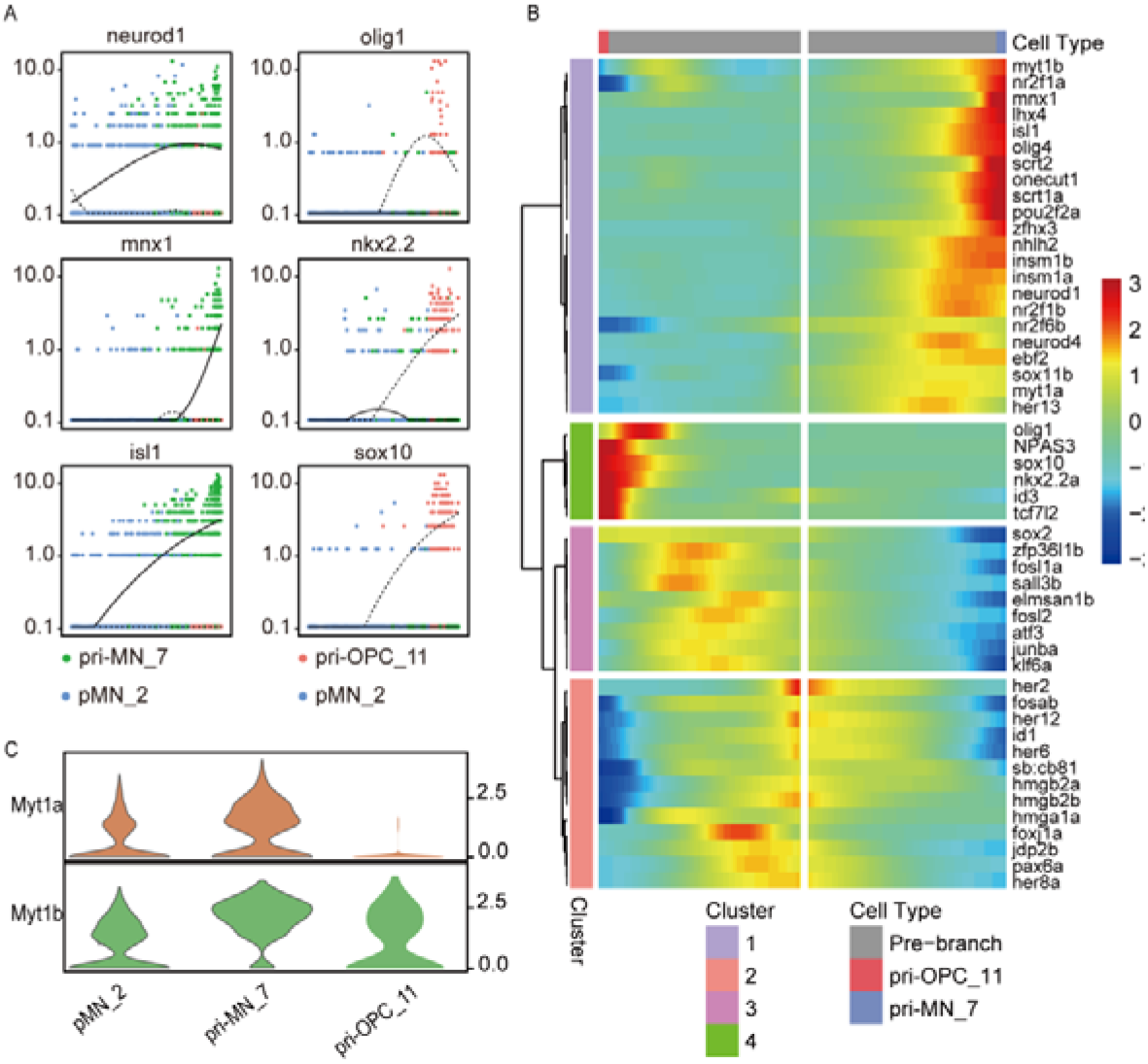
Transcription factor analysis for motor neuron and OPC lineages. A) expression of known markers in the branching of the motor neuron and OPC lineages. B) Heatmaps of the most differentially expressed transcription factors pre-branching, in the motor neuron and OPC lineages. The color red denotes upregulation; blue denotes downregulation of genes. C) Relative expression of *myt1a* and *myt1b* in the pMN progenitors and precursors shown by violin plots.

We observed that four distinct gene expression patterns along the MN or OPC trajectories: 1) a pattern of an increase in the MN trajectory (Cluster 1); 2) a pattern of an increase in the OPC trajectory (Cluster 2); 3) a pattern of a decrease in the MN but not in the OPC trajectory (Cluster 3); 4) a pattern of a decrease in either the MN or the OPC trajectory (Cluster 4). In Cluster 1, multiple transcription factors known for motor neuron specification, including *mnx1*, *lhx4*, *isl1*, *insml1a*, *insm1b*, and *neurod1*, were identified and defined as “MN transcription factors” (Figure 5A, B). In Cluster 2, transcription factors such as *olig1*, *sox10*, and *nkx2.2a* were included, consistent with their role in OPC lineage determination (Figure 5A, B). Cluster 3 and Cluster 4 included *sox2*, *hes6*, and *hes2*, which are known for counteracting cell differentiation programming and maintaining the stemness of neural stem cells or progenitors. Interestingly, genes previously identified as “regeneration-associated genes”, such as *sox11*, *atf3*, and *junb*, were distributed in either Cluster 1 or Cluster 3. This indicates that a complex dedifferentiation and neuron reprogramming process occurs in regeneration. These results were largely consistent with the DEG analysis between pri-MNs and pri-OPCs which revealed multiple differentially expressed genes or transcription factors for motor neuron or OPC fate determination (Supplemental Figure 6A-B), indicating the reliability of the different algorithms.

### The role of *myt1* in the neuronal lineage

We noticed that the transcription factors *myt1a* and *myt1b* (two isoforms of *myt1* in zebrafish) stood out in Cluster 1 (Figure 5B-C). *Myt1b* was significantly increased in the MN trajectory, while *myt1a* was increased in the MN trajectory but decreased in the OPC lineage. Myt1, myelin transcription factor 1, is known for its role in oligodendroctye differentiation (Nielsen, Berndt, Hudson, and Armstrong 2004), but its role in spinal cord circuit formation has not been well characterized.

To determine the role of *myt1* in motor neuron specification, we used CRISPR/Cas9-mediated gene disruption of *myt1a* and *myt1b* (Supplemental Figure 7A-B). We co-injected *myt1a* and *myt1b* sgRNAs and Cas9 protein into one-cell stage embryo, and found up to 100% mutagenesis in either *myt1a* or *myt1b* DNA amplicons of F0 embryos (n=11 and 13, respectively), as shown by Sanger Sequencing at 48 hpf (Supplemental Figure 7C-D). To test the extent of mutagenesis, we randomly selected three F0 samples and cloned 11 and 13 individual PCR amplicons in *myt1a* and *myt1b*, respectively. All clones had mutations (deletions, insertions or nucleotide substitutions), and many of them led to an out-of-frame protein product (Figure 6A-B, Supplemental 7E-F). Testing the stability of CRISPR, we found that the mutagenesis perdured for at least three months (Supplemental Figure 7G).

**Figure 6.**
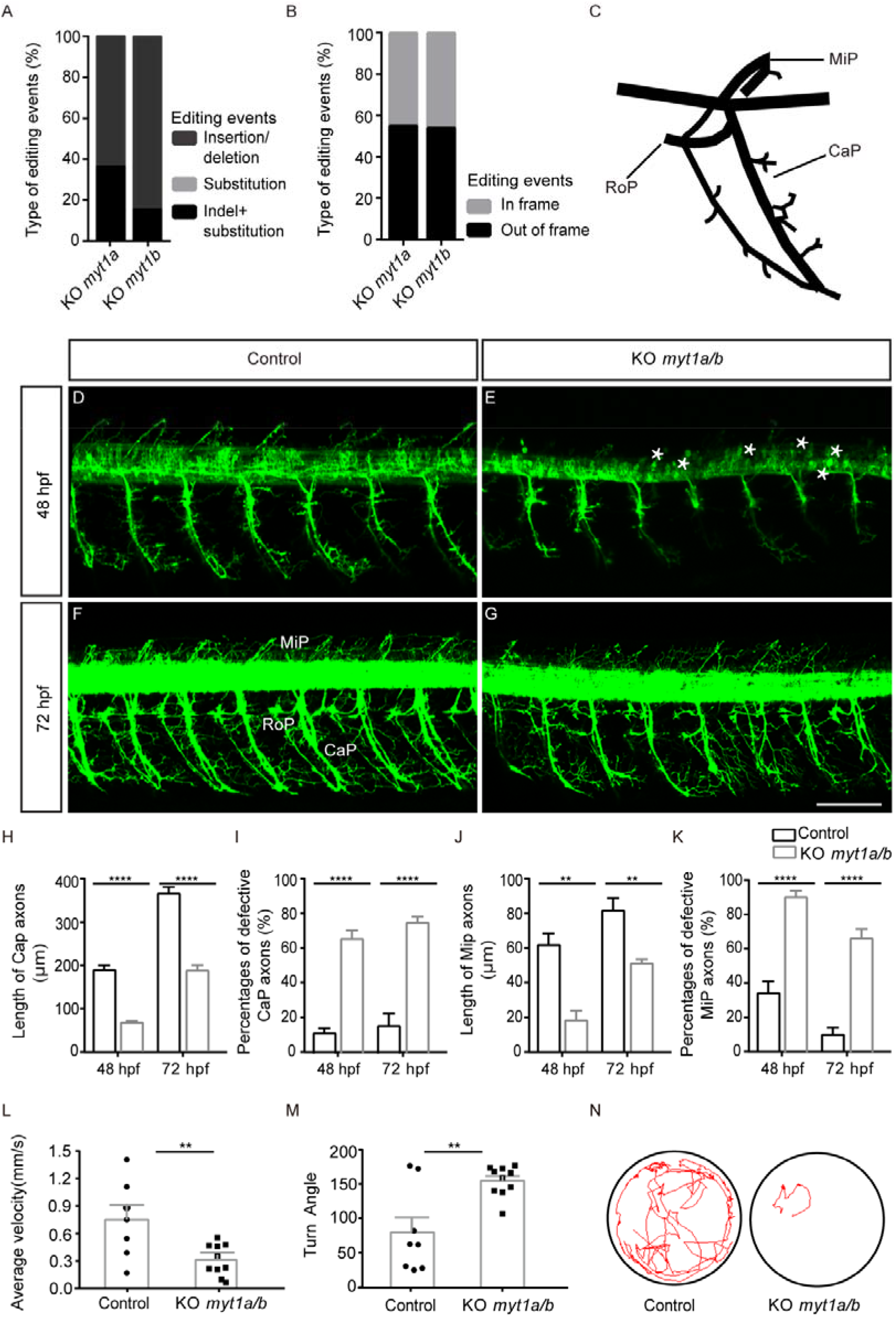
*Myt1* is necessary for the development of motor neurons. A-B) Types of *myt1a* or *myt1b* somatic mutagenesis generated by CRISPR. C) Schematic diagram of three primary motor axon projections (CaP, MiP and RoP) in zebrafish. D-G) Confocal imaging analysis of primary motor neurons in control and KO *Myt1a/b* groups at 48 hpf (D-E)and 72 hpf (F-G) Tg(hb9:GFP). Asterisks denote round-shaped and undifferentiated neuronal cells. Scale bar =50 µm. H) The length of CaP axons of zebrafish in the control group and in the KO *Myt1a/b* group at 48 hpf (n=5 and 5, respectively, t8=10.23, p<0.0001) and 72 hpf (n=5 and 5, respectively, t8=9.466, p<0.0001). I) The percentages of defective CaP axons in the control group and the KO *Myt1a/b* group at 48 hpf (n=5 and 7, respectively, t10=8.253, p<0.0001) and 72 hpf (n=5 and 7, respectively, t10=8.094, p<0.0001). J) The length of MiP axons of zebrafish in the control group and in the KO *Myt1a/b* group at 48 hpf (n=5 and 5, respectively, t8=4.968, p=0.0011) and 72 hpf (n=5 and 5, respectively, t8=3.957, p=0.0042). K) The percentages of defective MiP axons in the control group and the KO *Myt1a/b* group at 48 hpf (n=5 and 8, respectively, t11=7.602, p<0.0001) and 72 hpf (n=5 and 8, respectively, t11=7.259, p<0.0001). L) Velocity analysis in the control group and the KO *myt1a/b* group in 5 dpf larvae (n=7 and 10, respectively, t15=2.951, p=0.0099) by the video tracking software EthoVision XT. M) Turn angle in the control group and in the *myt1a/b* knockout group at 5 dpf larvae (n=8 and 10, respectively, t16=3.591, p=0.0024). *, **, and *** denote P < 0.05, P < 0.01, and P < 0.001, respectively. N) Representative locomotor trajectory of the control and the KO *myt1a/b*.

To analyze the development of motor neurons, we used the transgenic line *Tg(mnx1:GFP)* in which *GFP* was driven by the motor-neuron specific transcription factor *mnx1*. At 48 hpf when many motor neurons have been actively generated, most motor neurons in *myt1* mutant had round cell bodies and lacked processes, which was considered as undifferentiated states (Gong, Hu, Huang, Hu, Wang, Zhao, Qian, Wang, Sheng, Lu, Wei, and Liu 2020) (Figure 6E). In zebrafish, three primary motor neurons exist in the spinal cord: caudal primary motor neurons (CaP MNs), middle primary motor neurons (MiP MNs), and rostral primary motor neurons (RoP MNs), which innervate the ventral trunk musculature, dorsal trunk musculature, and muscle fibers in between, respectively. We viewed that many CaP or MiP axons were missing or shortened (Figure 6E). The average length of Cap axons were significantly shortened (n=5 embryo each, t8=10.23, p<0.0001; Figure 6H), and the defective Cap axons were dramatically increased in *myt1* mutant zebrafish (n=5 and 7, respectively, t10=8.094, p=0.0011; Figure 6I). The average length of MiP axons in mutants was only 1/3 of that in WT (n=5 embryo each, t8=4.968, p=0.0011; Figure 6J), and ∼90% MiP axons were missing or shortened in mutants (n=5 and 8 respectively, t11=7.602, p<0.0001; Figure 6K). Most RoP axons in the mutant were observable. However, it was difficult to determine their start and end points, which precluded further quantitative analysis (Figure 6G).

To exclude the possibility of developmental delay induced by CRISPR injection, we verified motor neuron specification at 72 hpf, a time point when motor neuron generation has been accomplished. In *myt1* mutants, the average length of Cap axons were only half of that in WT (n=5 embryo each, t8=9.466, p<0.0001; Figure 6H), and ∼80% of CaP axons had defects in their anterior lateral projections (n=5 and 7, respectively, t10=8.094, p<0.0001; Figure 6I). Similarly, MiP axons were still significantly shortened in mutants at 72hpf (n=5 embryo each, t8=3.957, p=0.0042; Figure 6J), and up to 60% MiP axons were missing or shortened (n=5 and 8, respectively, t11=7.259, p<0.0001; Figure 6K).

We also tested the behavioral outcomes of *myt1* mutagenesis in 5 day larvae. *Myt1* mutants have disrupted locomotor activity, as shown by a reduction in velocity (n=7 and 10 respectively, t15=2.951, p=0.0099; Figure 6L, N). Interestingly, we also found an increase in turn angle which was defined as the orientation of the head relative to the body when turning (n=8 and 10, respectively, t16=3.591, p=0.0024; Figure 6M). These indicate that both locomotion and motor coordination were disrupted in *myt1* mutants, validating *myt1*’s role in motor neuron development.

### The role of *npas3* in the development of the OPC lineage

We hypothesize that *npas3* may play a role in OPC lineage specification based on the fact that its expression is significantly increased in OPC precursors (Figure 5B). To test the role of *npas3* in OPC lineage specification, we designed a sgRNA targeted to exon 3 of *npas3*. Co-injection of *npas3* sgRNA and Cas9 led to a high rate of mutagenesis in F0 embryos, up to 100%. Sanger sequencing of single clone DNA samples of *npas3* crispants (n=12 clones) revealed that 100% contained frameshift mutations, leading to early truncation of the NPAS3 protein (Figure 7C, D, Supplemental Figure 8A-D).

**Figure 7.**
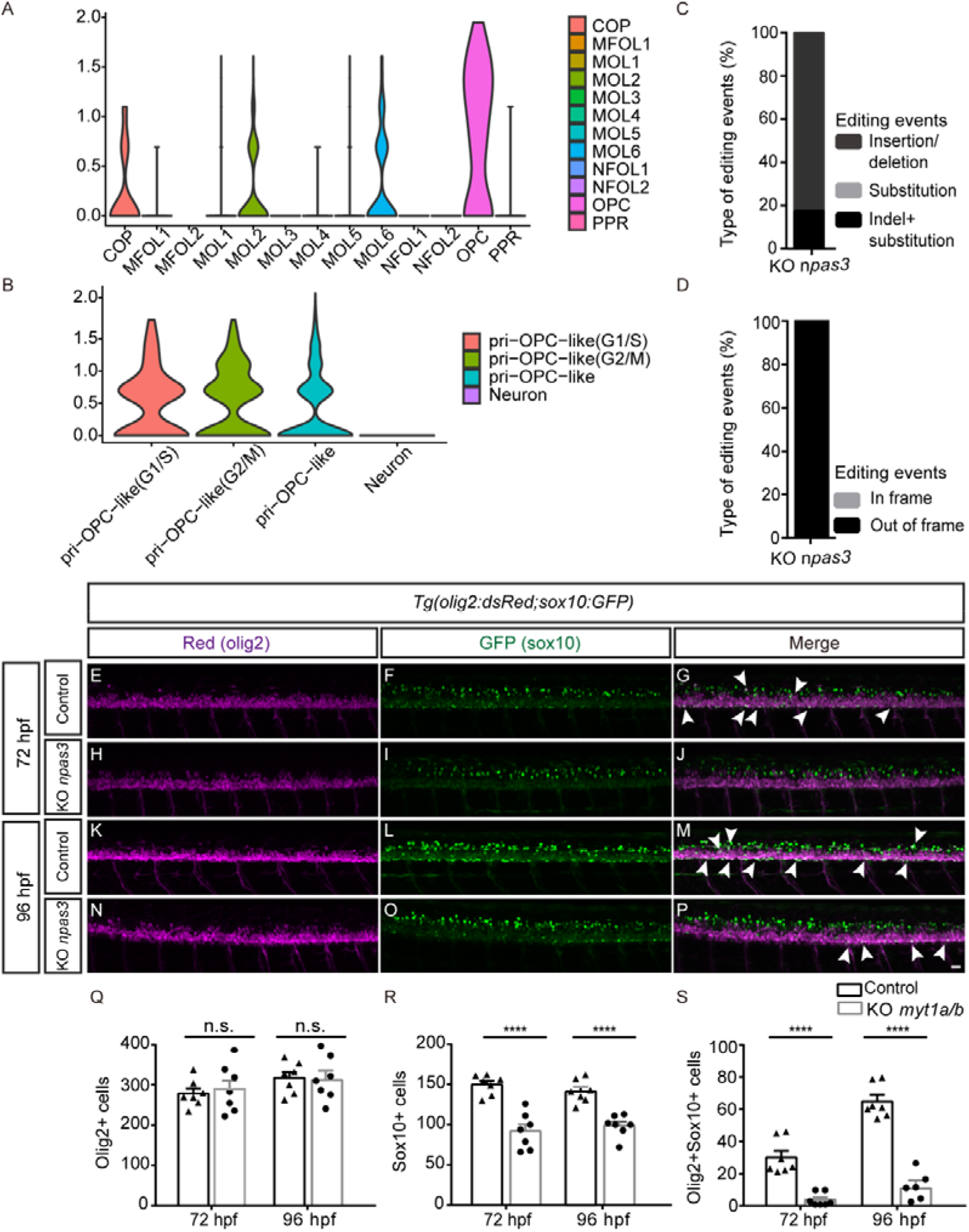
*Npas3* is necessary for the development of the OPC lineage. A-B) Violin plots of *npas3* in representative cells of the mammalian central nervous system from the datasets GSE128525 (A) and GSE122871 (B). Cells in GSE128525 were isolated from developing mouse spinal cords. Cells in GSE122871 were isolated from hGFAP:GFP mouse brain by FACS. C-D) Mutagenesis types generated by CRISPR. E-P) The OPC lineage defects observed in the *npas3* mutant at 72 hpf (E-J) and 96 hpf (K-P) in *Tg(sox10:GFP;olig2:dsRed)*. Scale bar=50 µm. Q) Quantification of *olig2*^+^ cells in controls and KO *myt1a/b*. (n=7 and 7, respectively, t12=0.4425, p=0.6660; n=7 and 7, respectively, t12=0.1013, p=0.9210. R) Quantification of *sox10*^+^ cells in controls and KO *myt1a/b*. (n=7 and 7, respectively, t12=5.974, p < 0.0001;n=7 and 7, respectively, t12=5.709, p < 0.0001) S) Quantification of *sox10*^+^*olig2*^+^ cells in controls and KO *myt1a/b*. (n=7 and 7, respectively, t12=6.126, p < 0.0001; n=7 and 6, respectively, t11=9.668, p < 0.0001). *, **, and *** denote p < 0.05, p < 0.01, and p < 0.001, respectively.

OPC lineages, including pri-OPCs, OPCs, and oligodendrocytes, are defined as cells labeled by *sox10* in the pMN domain (Scott, O’Rourke, Gillen, and Appel 2020). We analyzed OPC lineage with the transgenic line *Tg(sox10:GFP;olig2:dsRed)* at 72 hpf when the OPCs start to appear, and found that the number of *sox10*^+^ cells was reduced by half in the *npas3* mutant (n=7 embryo each, t12=5.974, p<0.0001; Figure 7R). This decrease was observed in both the ventral OPC and dorsal OPC lineage (Figure 7F, I). We observed that *olig2*^+^ cells were comparable in controls and *npas3* mutants (n=7 embryo each; t12=0.4425, p=0.6660; Figure 7Q; Figure 7E, H), indicating that this defect was not attributable to general developmental defects in the pMN domain, but rather was specific to the OPC lineage.

OPC lineage cells were also labeled by the marker *olig2*. To exclude the possibility of non-specific labeling of *sox10*-driven GFP in the transgenic line, we also analyzed the OPC lineage by counting the cells co-labeled with *sox10* and *olig2*. Consistently, the *npas3* mutant had a significant decrease in *sox10*^+^*olig2*^+^ cells (Figure 7G, J), with few double positive cells detected (n=7 embryo each, t12=6.126, p<0.0001; Figure 7S). Similar results were confirmed at 96 hpf, after OPCs have been specified. At 96 hpf, the number of *olig2*^+^ cells in the mutant group was almost the same as that in the control (Figure 7K, N; n=7 embryo each, t12=0.1013, p=0.9210 in Figure 7Q), but the *npas3* mutant still displayed a significant reduction in both *sox10*^+^ and *sox10*^+^*olig2*^+^ cells (Figure 7L-P; n=7 each embryo, t12=5.709, p<0.0001 in Figure 7R; n=7 and 6, respectively, t11=9.668, p<0.0001 in Figure 7S).

Notably, when we compared the expression of *npas3* in different cell clusters with two previously published single-cell datasets in mouse spinal cords and brains (Floriddia, Lourenco, Zhang, van Bruggen, Hilscher, Kukanja, Goncalves Dos Santos, Altinkok, Yokota, Llorens-Bobadilla, Mulinyawe, Graos, Sun, Frisen, Nilsson, and Castelo-Branco 2020; Weng, Wang, Wang, He, Cheng, Zhang, Verma, Xu, Dong, Liao, He, Potter, Zhang, Zhao, Xin, Zhou, Aronow, Blackshear, Rich, He, Zhou, Suva, Waclaw, Potter, Yu, and Lu 2019), we found that *npas3* was restricted in the OPC lineage (including pri-OPCs, OPCs and myelin) instead of neurons (Figure 7A-B), confirming a critical role of *npas3* in the OPC lineage development.

### The association between *npas3* and other schizophrenia genes

Disruption of *NPAS3* is associated with schizophrenia and learning disabilities (Pickard, Malloy, Porteous, Blackwood, and Muir 2005; Kamnasaran, Muir, Ferguson-Smith, and Cox 2003), and its role in neurogenesis has been identified (Pieper, Wu, Han, Estill, Dang, Wu, Reece-Fincanon, Dudley, Richardson, Brat, and McKnight 2005). However, how NPAS3 functions in neurodevelopmental and psychiatric disorders is largely uncharacterized. A comparison of the amino acid sequences of NPAS3 revealed that the entire protein in zebrafish is 74.3% and 75% identical to its human and mouse orthologs, respectively (Figure 8A, Supplemental Figure 9). NPAS3 encompasses two functional domains: the basic helix-loop-helix (bHLH) and PAS domains, which are critical for DNA and protein interaction (Luoma and Berry 2018). The basic helix-loop-helix domain in zebrafish is 98.2% identical to its human and mouse orthologs. The PAS domain is 96.6% identical to its human and mouse orthologs. Mouse and human NPAS3 share the same sequences in both the bHLH and PAS domains. The NPAS3 protein, especially in its functional domains, maintains a high degree of conservation.

**Figure 8.**
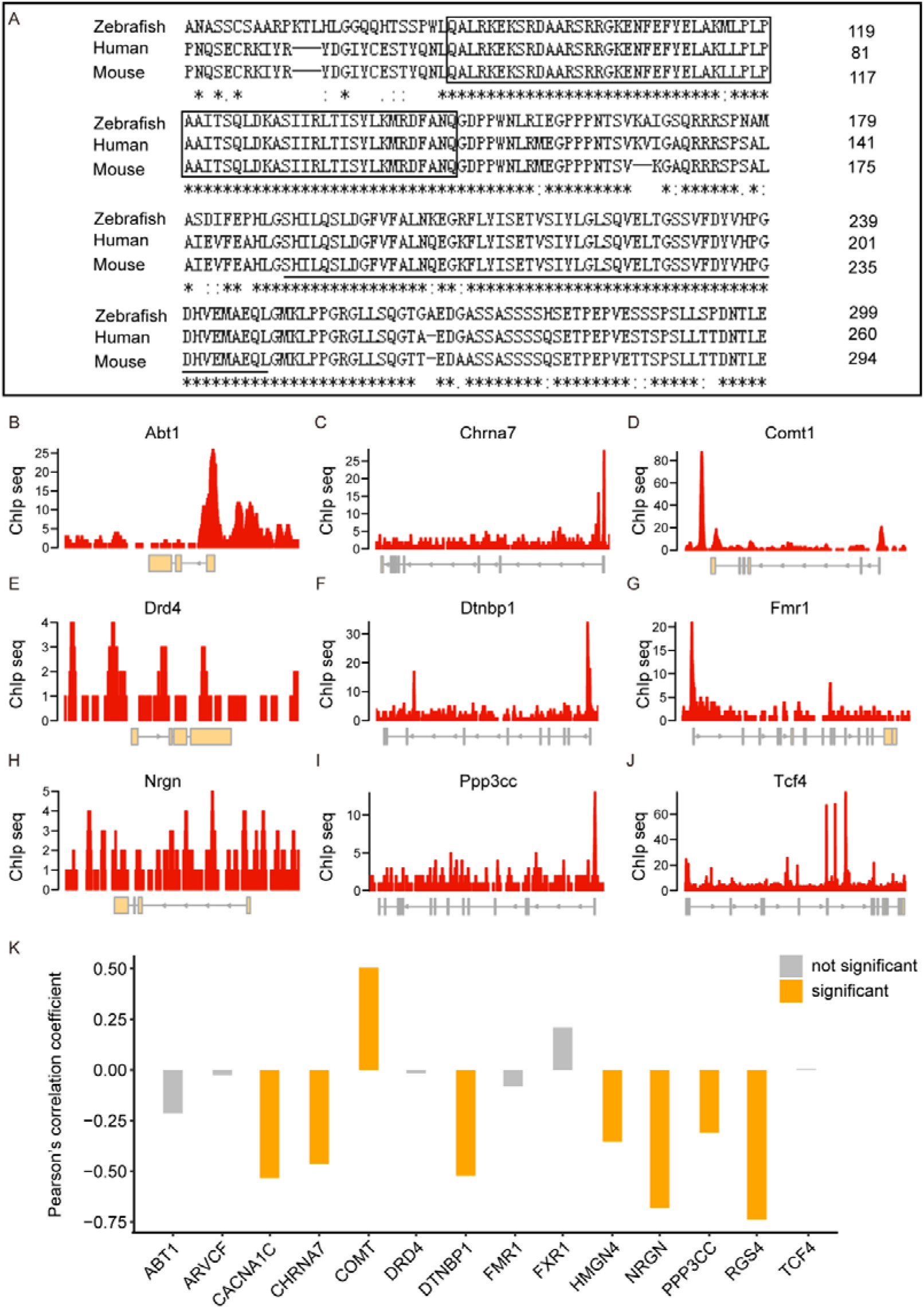
The association analysis of NPAS3 and other “Schizophrenia Genes”. A) Alignment of NPAS3 amino acids with the ClustalW. “*”depicts identical amino acids; “:” represents amino acids with strong similarity; “.” denotes a site with weak similarity. The bHLH domain is boxed, and the PAS domain is underlined. B-J) Binding site profile of *npas3* and other “Schizophrenia Genes”. Exons are denoted as boxes, and introns are represented as lines. Data was retrieved from GSE70872. K) A correlation analysis between *npas3* and other “Schizophrenia Genes” based on their gene expression levels. The expression values were calculated according to the log (RPKM) in the transcriptome data (GSE78936). (RPKM: Reads Per Kilobase per Million mapped reads). The Pearson correlation coefficient thresholds >0.25 or <-0.25 were considered as significant associations (shown as an orange box).

Schizophrenia is a complex neuropsychiatric disorder which can be attributed to both genetic and environmental risk factors. Twin and adoption studies reveal a high degree of heritability, and many genes have been found to be associated with schizophrenia, including NPAS3 (Gejman, Sanders, and Duan 2010; Pickard, Malloy, Porteous, Blackwood, and Muir 2005). To determine the potential role of NPAS3 in schizophrenia, we used NPAS3 ChIP (chromatin immunoprecipitation) data to analyze the physical interactions between NPAS3 and genes reported to be associated with schizophrenia (designated “Schizophrenia Genes”) (Gejman, Sanders, and Duan 2010; Thyme, Pieper, Li, Pandey, Wang, Morris, Sha, Choi, Herrera, Soucy, Zimmerman, Randlett, Greenwood, McCarroll, and Schier 2019). These “Schizophrenia Genes” we selected were found to be associated with schizophrenia in at least two independent studies (Supplemental Table 2). These include COMT, DTNBP1, and NRG1, the most studied genes related to schizophrenia (Henriksen, Nordgaard, and Jansson 2017). This mouse ChIP data (Moen, Adams, Brandsma, Dekkers, Akinci, Karkampouna, Quevedo, Kockx, Ozgur, van, Demmers, and Poot 2017) (human data is not available) is a powerful approach to discover the downstream targets of NPAS3, since both human and mouse NPAS3 have exactly the same sequences in both functional domains (Figure 8A, Supplemental Figure 9).

In this dataset, 19 out of 31 “Schizophrenia Genes” were found. NPAS3 can bind to the promoter regions in eight of these genes, including *abt1*, *chrna7*, *dtnbp1*, *fmr1*, *ppp3cc*, *comt1*, *cacna1c* and *tcf4*. NPAS3 can bind to the enhancer regions in six of these genes, including *arv4cf*, *drd4*, *fxr1*, *nrgn*, *hmgn4*, and *rgs4*. These data indicate a potential interaction between NPAS3 and other “Schizophrenia Genes” (Figure 8B-J, Supplemental Figure 10A-E).

We retrieved high-throughput RNA-seq data in patients with schizophrenia and their healthy controls (Hu, Xu, Pang, Zhao, Li, Deng, Liu, Lan, Zhang, Zhao, Xu, Xu, Xiao, and Li 2016), and tested the correlation between the *npas3* gene and “Schizophrenia Genes” based on their expression levels using Pearson’s correlation. After filtering genes with low expression levels (RPKM<1), all of these 14 “Schizophrenia genes” bound by NPAS3 in their promoter regions or enhancer regions were detected. We found that *npas3* was negatively associated with most of these genes, including *chrna7, dtnbp1, ppp3cc, cacna1c, nrgn, hmgn4,* and *rgs4*. *Comt*, however, was positively associated with *npas3* (Figure 8K).

We have identified that *npas3* can bind to a number of “Schizophrenia genes”, and *npas3* was significantly correlated with the expression levels of many of these genes. These offer potential roles of this transcription factor in schizophrenia pathogenesis.

### Web-based tools for exploration

Cell cluster definition is not well established in zebrafish due to the relative paucity of databases, and zebrafish have unique cellular expression signatures compared to mammals. We generated a web-based tool for our scRNA-seq datasets, including not only *olig2*^+^ cells (progenitors, precursors, as well as multiple states of motor neurons) but also interneurons and other cells sorted by FACS. With this interface, users can visualize gene transcription levels and patterns and/or explore highly expressed genes in each individual cluster (Figure 9A-D). This tool is available at: https://nantongneurokeylab.shinyapps.io/cell_browser/.

**Figure 9.**
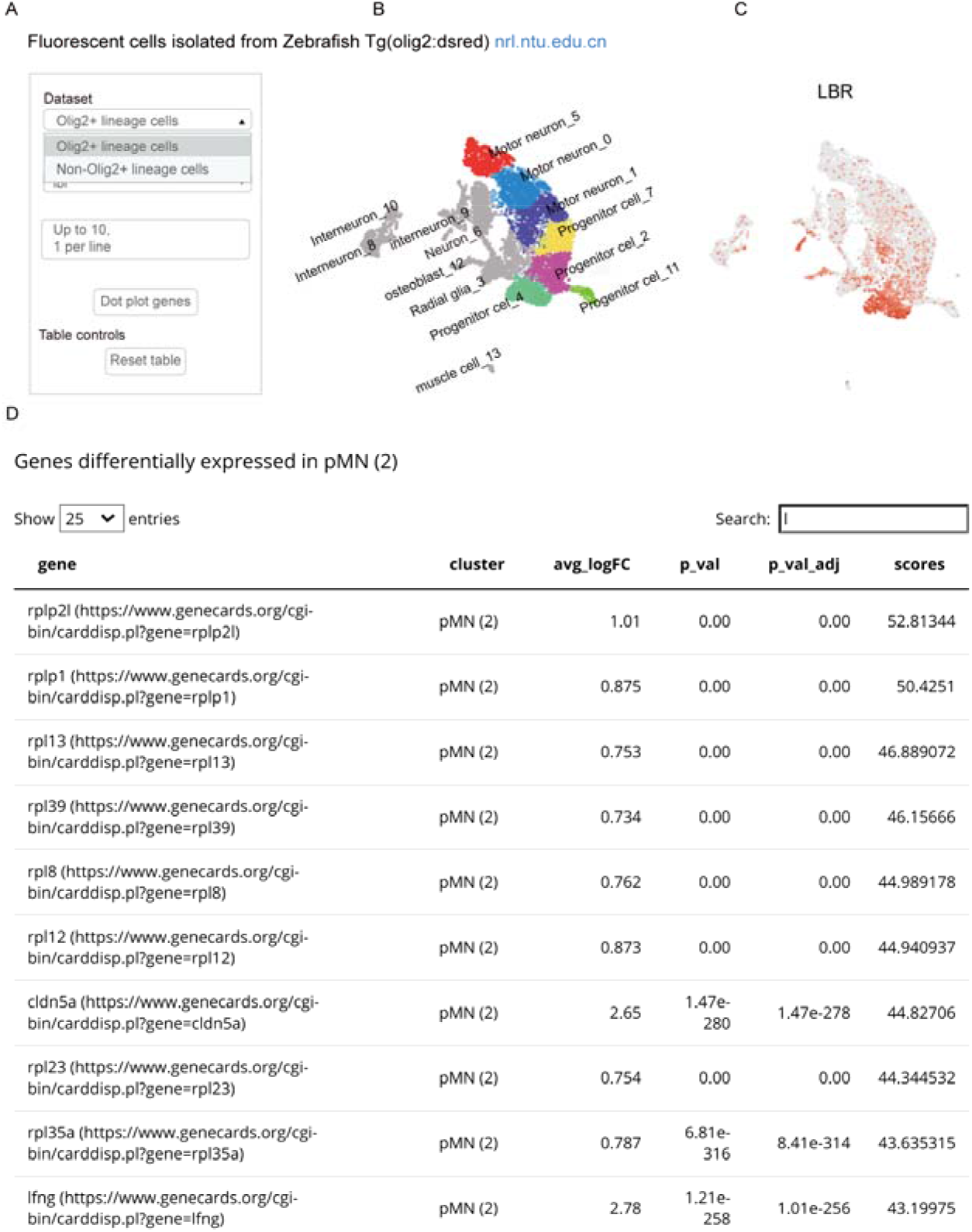
Web-based single cell datasets. A) Gene search window. B) UMAP visualization of cell clustering. C) Expression of genes searched shown with UMAP. D) Top differentially expressed genes in the cell subpopulation searched.

## Discussion

Our study, utilizing single-cell RNA sequencing, has identified multiple types of pMN progenitors/precursors, including proliferating progenitors, differentiating progenitors, and restricted precursors for neurons and OPCs. Lineage analyses by Monocle or PAGA revealed that both motor neurons and OPC precursors are derived from the same pMN progenitors but distinct lineage-restricted precursors. We also identified some of the molecular programs for progenitor fate determination, showing that *myt1* and *npas3* are necessary for motor neuron and OPC lineage specification, respectively (Figure 10). Our scRNA-seq has not only deciphered the lineages of pMN progenitors, but has also identified regulators of cell fate determination.

**Figure 10.**
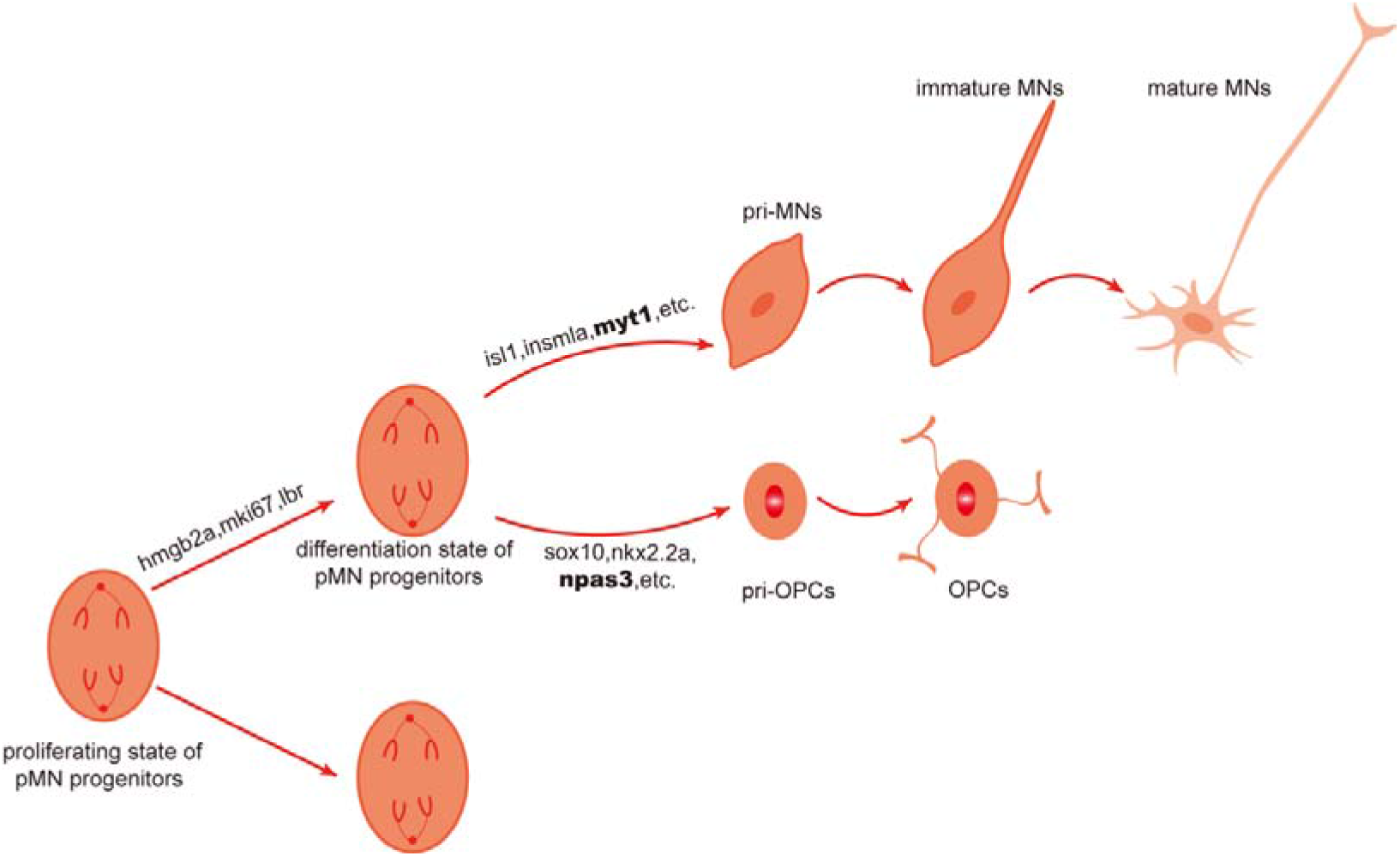
Diagrams of olig2+ cell lineages and molecular regulators for cell fate determination. OPCs and motor neurons derive from common olig2+ pMN progenitors but with distinct restricted precursors.

Zebrafish cells were sorted at 42 hpf when motor neurons are actively produced and cells from the OPC lineage begin to appear. Though it is likely that additional scarce pMN progenitors remained undetected, cell sorting revealed the existence of precursors for motor neurons and OPCs, and that the precursors are derived from the same *olig2*-expressing progenitors. This is consistent with the finding that the progeny of clones isolated from neural tissue in mice, chick, and zebrafish contain lineage cells for both motor neurons and OPCs (Leber, Breedlove, and Sanes 1990; Leber and Sanes 1995; Park, Shin, and Appel 2004). For this “Common Progenitors” model, one possibility is the “Switching” model in which pMN progenitors first generate motor neuron precursors then switch to generating OPCs; the other is the “Segregating” model in which motor neurons and oligodendrocyte precursors segregate early but the OPC specification initiates only when motor neuron generation ceases (Richardson, Smith, Sun, Pringle, Hall, and Woodruff 2000). At 42 hpf when motor neurons are still actively generated, we observed OPC precursors instead of OPC (high expression of *sox10* and *nkx2.2* but low expression of *pdgf*α) along with motor neuron precursors, which supports the “Segregating” model. It is likely that in early development pMN progenitors stochastically develop into motor neuron or oligodendrocyte lineage fates. This scenario may follow the same rule as observed in drosophila in which neuroblast express a sequence of transcription factors specify progeny cells when dividing (Isshiki, Pearson, Holbrook, and Doe 2001; Cleary and Doe 2006; Kao, Yu, He, Kao, and Lee 2012; Bayraktar and Doe 2013). Spinal cords and brains have distinct progenitor pools (Metzis, Steinhauser, Pakanavicius, Gouti, Stamataki, Ivanovitch, Watson, Rayon, Mousavy Gharavy, Lovell-Badge, Luscombe, and Briscoe 2018), but progenitors and their offspring in spinal cords or brains share similar markers. To exclude the possibility of confusing lineages, our study focused only on progenitors in the spinal cords. It would be interesting to decipher whether the progenitor lineage trajectories in the brains follow the same rule.

Interestingly, the diversity of pMN progenitors suggests a new interpretation for the apparent conflict of the motor neuron and OPC lineages. Our single cell diversity is consistent not only with the evidence for the segregating model, but also with the evidence found for the distinct progenitors model, in which motor neurons derived from pMN progenitors seldom divide (Ravanelli and Appel 2015). As we show, both pri-MN and pri-OPC derived from the same subpopulation pMN_2, the differentiating state of pMN progenitors (Figure 2A, Figure 4C-D). We found that the transcription factor *olig2* is enriched in pri-MN, pri-OPC, and their progenitors. This may explain why previous results have misleading conclusions for the lineages of cells traced by this single *olig2* reporter (Park, Shin, and Appel 2004; Ravanelli and Appel 2015), depending on which specific type of cells have been selected. Our study shows that with only a single marker/reporter, *olig2,* for example, it is hard to distinguish different subpopulations of progenitors, and this may lead to inaccurate lineage tracing. Our interactive website offering signature genes in each subpopulation will give researchers access to a powerful tool to develop more specific markers for cell labeling. Zebrafish, rodents, and humans share common genes but also possess specific genes in cell subpopulations. Therefore, this website may also offer a robust tool for comparative biology.

Lineage analysis of the single-cell transcriptome deepens our understanding of molecular programs driving neuron or OPC fate determination. The divergence of pri-MN and pri-OPC from the common olig2+ progenitors indicates that molecular events are critical for fate transition. Consistently, many genes enriched in pri-OPC or pri-MN lineages have known roles in motor neuron or OPC specification, respectively. The validation of the role of myt1 and npas3 in the specification of motor neurons or OPCs confirms that lineage deciphering is a powerful tool for molecular programming analysis. In spite of the absence of OPC cells in our isolated cells, multiple markers for pri-OPCs indicate that these pri-OPCs will become OPCs. It would be interesting for future studies to sort cells at later stages and dissect other possible fates of pri-OPCs, e.g. the transition into astrocytes.

Early studies focused on the role of myt1 in oligodendrocyte differentiation (Nielsen, Berndt, Hudson, and Armstrong 2004), and its role in neuron specification has been recognized recently (Vasconcelos, Sessa, Laranjeira, Raposo, Teixeira, Hagey, Tomaz, Muhr, Broccoli, and Castro 2016). Our work shows that *myt1* is specifically upregulated in motor neuron precursors rather than OPC precursors, indicating that it may function in glia-neuron fate determination. It may be interesting to explore how this transcription factor works with general neuronal transcription factors, e.g. Neurogenin-1 and Neurogenin-2, and motor neuron-specific transcription factor, e.g. *insm1* or *insm2*. In addition, we noticed that *myt1* is specifically expressed in neuronal cells, including the motor neuron lineage and interneurons (see our interactive website), indicating its critical role in neuronal fate determination. The Myt1 family includes three proteins: Myt1, Nzf3, and Myt1l (Myt1-like). Myt1l has been implicated in neuronal reprogramming either alone or in combination with the transcription factors *ascl1* and *bcl2* (Vierbuchen, Ostermeier, Pang, Kokubu, Sudhof, and Wernig 2010; Liang, Tan, Wang, Tao, Huang, Ma, Fukunaga, Huang, and Han 2018). Neuronal reprogramming has been applied to generate neurons in many diseases and pathological states, including spinal cord regeneration, Parkinson’s Disease, and Alzheimer’s Disease (Yang, Xing, Yu, cai, Cui, and Chen 2020; Guo, Zhang, Wu, Chen, Wang, and Chen 2014; Qian, Kang, Hu, Zhang, Liang, Meng, Zhang, Xue, Maimon, Dowdy, Devaraj, Zhou, Mobley, Cleveland, and Fu 2020). Therefore, it might be interesting to explore the role of Myt1 in neuronal reprogramming, which may offer new insight into neuronal regeneration.

*Npas3* insufficiency is associated with schizophrenia and mental retardation (Pickard, Malloy, Porteous, Blackwood, and Muir 2005; Kamnasaran, Muir, Ferguson-Smith, and Cox 2003). In parallel, *npas3* knockout mice have provocative behavior and altered anxiety-related responses (Brunskill, Ehrman, Williams, Klanke, Hammer, Schaefer, Sah, Dorn, Potter, and Vorhees 2005). Deficits in hippocampal neurogenesis have also been observed in *npas3* knockouts (Pieper, Wu, Han, Estill, Dang, Wu, Reece-Fincanon, Dudley, Richardson, Brat, and McKnight 2005). In fact, oligodendrocyte differentiation and myelin defects contribute to schizophrenia and learning disabilities as well as neuronal defects (Takahashi, Sakurai, Davis, and Buxbaum 2011; Terzi, Oğuzkan-Balci, Anlar, Erdoğan-Bakar, and Ayter 2011). Our result that NPAS3 is necessary for OPC specification may provide a novel role for NPAS3 in psychiatric disorders. Surprisingly, the downstream signaling of NPAS3 is largely uncharacterized. Our ChIP and transcriptome data analysis suggest that NPAS3 may have interaction with genes associated with schizophrenia, many of which have significant role in neural circuit formation. Functional tests are necessary to further confirm the downstream signaling of NPAS3 in human OPC lineage cells.

In conclusion, our work provides insights into lineages of neurons and glia, which is essential for understanding the basis of nervous system circuit formation. The identification of critical programming in cell fate determination may also offer clues for understanding central nervous system disorders and may be exploited for neuronal regeneration.

## Methods

### Ethics Statement

All animal experiments were performed in accordance with the NIH Guide for the Care and Use of Laboratory Animals (http://oacu.od.nih.gov/regs/index.htm). All procedures and protocols were approved by the Administrative Committee for Experimental Animals, Jiangsu Province, China (Approval ID: SYXK (SU)2007–0021).

### Zebrafish husbandry and collection of embryos

Wild type and transgenic zebrafish *Tg(olig2:GFP)^vu1^* (Shin, Park, Topczewska, Mawdsley, and Appel 2003), *Tg(sox10*:*GFP)^ba2^* (Carney, Dutton, Greenhill, Delfino-Machin, Dufourcq, Blader, and Kelsh 2006), and *Tg(mnx1*:*GFP)^ml2^ (Flanagan-Steet, Fox, Meyer, and Sanes 2005)* were obtained from the China Zebrafish Resource Center (CZRC). Zebrafish feeding and embryo manipulations were conducted according to standard protocols in the Nantong Zebrafish Core Facility. The adults were raised at 28 °C, with 14/10 hour light-dark cycles. Embryos were raised in petri dishes at 28 °C.

### Single cell isolation and sorting

The transgenic line *Tg (olig2:dsRed)* was outcrossed with wild type, and their embryos were anesthetized at 42 hour post fertilization (hpf) with 0.04% MS-222 and dechorionated. Heads were removed rostral to the hindbrain-spinal cord boundary using a 24 G needle. The remaining trunks were deyolked in deyolking buffer (55□mM NaCl, 1.8□mM KCl, and 1.25□mM NaHCO_3_) and minced with fine scissors. Samples were dissociated with 0.25% trypsin at 28°C for 30 minutes followed with 1 mg/ml collagenase II for 20 minutes. 10% FBS was added to terminate the enzymatic reaction. Samples were passed through a 70 um strainer and sorted with a BD FACS-Aria fusion machine as previously reported (Weinberger, Simoes, Patient, Sauka-Spengler, and Riley 2020).

### Single cell capture, sequencing, and data processing

The cell suspension (300-600 cells/ul) was loaded onto a 10X Chromium platform according to the manufacturer’s instructions. Captured cells were lysed, and released RNAs were reverse transcribed into cDNA. cDNAs were amplified and libraries were generated with the Single Cell 3’ Library and Gel Bead kit V3. The libraries were sequenced on an Illumina Novaseq 6000 with at least 100,000 reads per cell (CapitalBio Technology, Beijing) using 150 bp paired-end runs. The Cell Ranger, downloaded from https://support.10xgenomics.com/single-cell-gene-expression/software/downloads/latest, was used to de-multiplex Illumina output and generate feature-barcode matrices.

### Cell clustering and lineage analysis

Genes detectable in more than two cells were defined as “expressed”. Cells with <1000 genes detected were removed. Counts were normalized to 10000 and transformed into logarithmic scales. A neighborhood graph was embedded using UMAP displaying the top 4000 highly variable genes across cells. Cells were clustered by the Louvain Algorithm. The trajectory interface of all cell clusters was done using the layout ‘fa’. These analyses were performed by the Python-based SCANPY package, designated as partition-based graph abstraction (PAGA) (Wolf, Angerer, and Theis 2018). Trajectory lineages of pMN progenitors or precursors were also analyzed with Monocle with built-in R (Trapnell, Cacchiarelli, Grimsby, Pokharel, Li, Morse, Lennon, Livak, Mikkelsen, and Rinn 2014). For PAGA analysis, clustering of all cells as well as clustering of subpopulations of pMN progenitors and precursors were performed. For Monocle analysis, only subpopulations of pMN progenitors/precursors were analyzed.

### *Npas3* or *myt1* CRISPR/Cas9

One-cell stage zebrafish embryos from *Tg(mnx1:GFP)^ml2^* (Flanagan-Steet, Fox, Meyer, and Sanes 2005) outcrossed to wild type were injected with 450 pg Cas9 protein (Novoprotein, E365), 225 pg *myt1a* and 225 pg *myt1b* sgRNAs. SgRNA synthesis was performed as previously reported (Ding, Gu, Cai, Cai, Yang, Bao, Shen, Ni, Chen, and Xing 2020). Briefly, double-stranded DNA for the gRNA was PCR amplified with the template vector pMD 19-T and the following primers were used: *myt1a* sgRNA forward: GGGGGACTCCTGGAACAGGC, *myt1b* sgRNA forward: GGCCGCAGTCTGTCAGGCTG, *npas3* sgRNA forward: GGGTCCCGGGAGCTACAAAC and universal reverse: AAAAAAAGCACCGACTCGGTGCCAC. Amplicons were purified with the DNA clean & concentrator kit (Zymo Research). Functional sgRNA was transcribed using the MegashortscriptTM kit (Thermo Fisher Scientific) and purified with the Megaclear kit (Thermo Fisher Scientific).

### Verification of mutagenesis generated by CRISPR

Genomic DNA was extracted from 48 or 72 hours post-fertilization (hpf) embryos and analyzed as previously described (Xing, Son, Stevenson, Lillesaar, Bally-Cuif, Dahl, and Bonkowsky 2015). In brief, 50 mM NaOH was used to lyse embryos heated at 95°C for 30 min. DNA surrounding the *npas3*, *myt1a*, and *myt1b* sgRNA regions was amplified with their respective primers: *npas3* F:GAGCTTGAAAGCTTGGAGCG R:GTTTTGCGGTTAGTCTGCGT; *myt1a* F:GACCACCAGTACTTCAGCGG R:CGGCGATGTCCATGACCTTA; *myt1b* F:AGCATGAAACTGGAGAGCGT R:TCGAAGGCACGGATTGTCTT. The targeted DNA regions were purified and sequenced by Sanger sequencing. To analyze the mutagenesis frequency, amplified PCR products from F0 embryos were cloned into PCR4.0 TOPO (ThermoFisher). Plasmids from individual colonies were isolated and sequenced by Sanger sequencing (Genewiz, Inc).

### Analysis of larval locomotor activity

Larval behavior was tracked at 5 dpf with an Ethovision XT 13.0 equipped with a Basler GenlCam (Basler acA 1300-60) at a rate of 25 images/sec. At 5 dpf, larvae were transferred into a 24-well plate with one larva per well. After acclimatization in the DanioVision Observation Chamber (Noldus Information Technology) for 5 minutes in the dark, swimming behavior was recorded for 10 minutes. We used a light/dark flash model with 10 s (seconds) light followed by 10s dark (thirty cycles, for a total of 10 min). This model was to evaluate the response of larvae to acute illumination changes. The light/dark program settings of this method were derived from the manufacturer’s manual. The parameters we analyzed included total distance moved, average speed, angular velocity, and turn angle. To calculate total distance moved, we inferred distance moved relative to the center. 0.2 millimeter (mm) was set as the threshold value to exclude slight movements that were not considered to be swimming.

### *In situ* probe generation and fluorescent in situ hybridization

*In situ* RNA probes for *olig2*, *mnx1*, *btg2*, *sox10*, and *hes6* were generated by a plasmid-free approach as previously (Xie, Kaufmann, Moulton, Panahi, Gaynes, Watters, Zhou, Xue, Fung, Levine, Letsou, Brennan, and Dorsky 2017). Primers used for probes were listed in Table 1. cDNAs from 2 and 3 dpf embryos were amplified with related primers, and PCR products were purified with Zymo Research DNA Clean & Concentrator-5 kit. T7 polymerase was used to make digoxigenin-, fluorescein- and DNP-labeled antisense probes for two-color or three-color *in situ* hybridization. Whole-mount in situ hybridization was performed as described in ZFIN (https://zfin.atlassian.net/wiki/spaces/prot/pages/365134287/Triple+Fluorescent+In+Situ).

### Microscopy and image analysis

Confocal images of transgenic zebrafish were acquired as previously described (Ding, Gu, Cai, Cai, Yang, Bao, Shen, Ni, Chen, and Xing 2020). Live *Tg(olig2:dsRed;sox10:GFP)* and *Tg(mnx1:GFP)* embryos were mounted lateral side up in low-melt agarose. Confocal z-stacks were taken with the same confocal settings (e.g. the same PMT and imaging speeds). Analysis was performed on max Z projections in ImageJ. ∼7 consecutive spinal cord segments in the confocal imaging windows were analzyed, with the most rostral segment over the caudal margin of the yolk sac. To quantify the length of the CaP axons, we imported the images into Imaris Software (Oxford Instruments), traced the axons, and calculated their lengths. In addition, we used ImageJ to calculate the length of RoP and Mip axons. For quantification of the OPC lineage cells, puncta were anazlyed with ‘analzye particles’ in ImageJ with a size of 0.01 to infinity and a circularity of 0.01 to 1.00.

### The association analysis of NPAS3 and other “Schizophrenia Genes”

The high-risk genes for schizophrenia were selected according to the literature (Gejman, Sanders, and Duan 2010; Thyme, Pieper, Li, Pandey, Wang, Morris, Sha, Choi, Herrera, Soucy, Zimmerman, Randlett, Greenwood, McCarroll, and Schier 2019). Since many genetic association studies have inconsistent results, we further queried for publications about each gene and schizophrenia in Pubmed. We manually reviewing the titles and abstracts of each paper. Genes were retained when at least two independent genetic association studies or genome-wide association analysis (GWAS) found a correlation with schizophrenia (Supplemental Table 2). These genes were defined as “Schizophrenia Genes”. The correlation analysis between NPAS3 and other “Schizophrenia Genes” was based on their gene expression levels in schizophrenia patients and their healthy controls (Hu, Xu, Pang, Zhao, Li, Deng, Liu, Lan, Zhang, Zhao, Xu, Xu, Xiao, and Li 2016). The GSE78936 datasets were high throughput RNA-seq data. After removing the outliers (BA11_17, BA11_22, BA11_23, BA11_25, BA11_26, BA9_13, BA24_12), 45 samples were left, which included 17 controls and 28 schizophrenic samples. After filtering genes with at least 30% samples with RPKM<1, 29 genes were left. The expression values were calculated by log (RPKM). (RPKM: Reads Per Kilobase per Million mapped reads). Pearson correlation coefficients between npas3 and other “Schizophrenia genes” were calculated by the R function cor.test.

To test the interaction between NPAS3 and other “Schizophrenia genes”, the published ChIP-seq data of NPAS3 were retrieved (Moen, Adams, Brandsma, Dekkers, Akinci, Karkampouna, Quevedo, Kockx, Ozgur, van, Demmers, and Poot 2017). The “Gviz” Package in R (Hahne and Ivanek 2016) was applied to visualize the peak information with default settings.

### Statistical analysis

GraphPad Prism (version 6.01) was used for statistical analysis of the data. For confocal image analysis, ten 3μm Z intervals of consecutive trunk spinal cord sections were stacked with the max Z projection in Image J. For analysis in two groups, a two-tailed t-test (unpaired) was performed. Data were shown as the mean ± standard error of the mean (SEM).

### Data Availability Statement

The data for this study are available in the Sequence Read Archive of the National Institutes of Health (SRA accession: PRJNA722570), and the interactive website https://nantongneurokeylab.shinyapps.io/cell_browser/. Computer codes are available at: https://github.com/xinglingyan/Single-Cell-Analysis.

## Acknowledgements

Funding:This work was funded by the National Key Research and Development Program of China (2017YFA0104704), the National Natural Science Foundation of China (81701127, 31872773, 32070998), the Key Research and Development Program (Social Development) of Jiangsu Province (BE2020667), the Foundation of Jiangsu Province “333 Project High-level Talents” (BRA2020076) and the Priority Academic Program Development of Jiangsu Higher Education Institutions (PAPD).

## Author contributions

LX: Investigation, data analysis, conceptualization, funding acquisition, and writing–original draft, review & editing; RC: Investigation and writing-original draft; JW: Investigation, data analysis, and writing-original draft. HL: data analysis; JL: Investigation; YW: Website dataset generation; BL: data analysis; JS: Data analysis; GC: Supervision and funding acquisition.

---

**Supplemental Figure 1:**
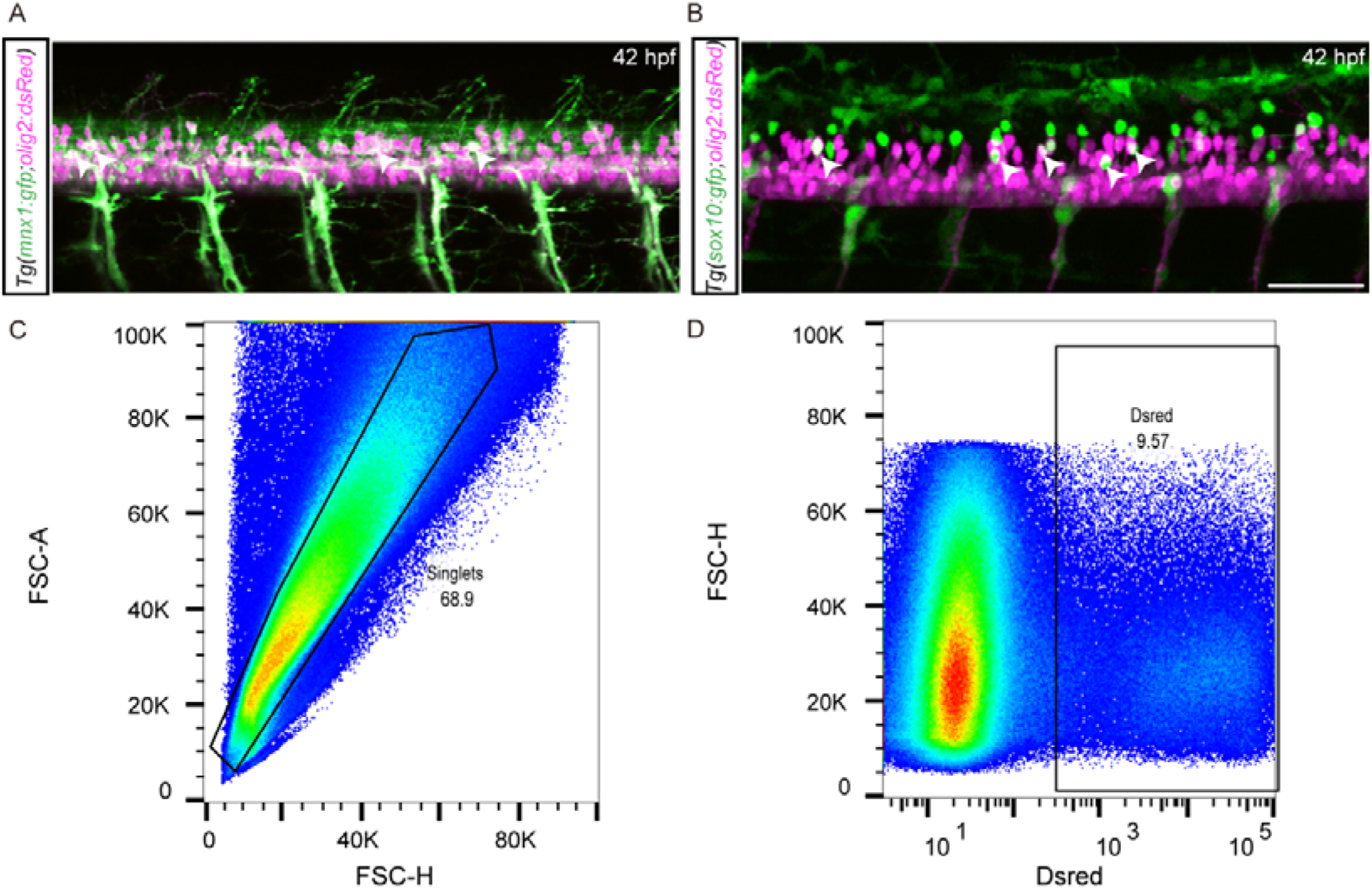
Sorting of olig2+ cells by FACS. A) *Tg(olig2:dsRed;mnx1:GFP)* embryo double-labeled for olig2 progenitors and motor neurons. B) *Tg(olig2:dsRed;sox10:GFP)* embryo double-labeled for olig2 progenitors and OPC lineage cells. A-B) Confocal images of whole-mount embryos at 42 hpf, lateral view. Scale bar, 50 μm. C-D) Sorting strategy for the Olig2+ cells from *Tg(olig2:dsRed)* by FACS.

**Supplemental Figure 2:**
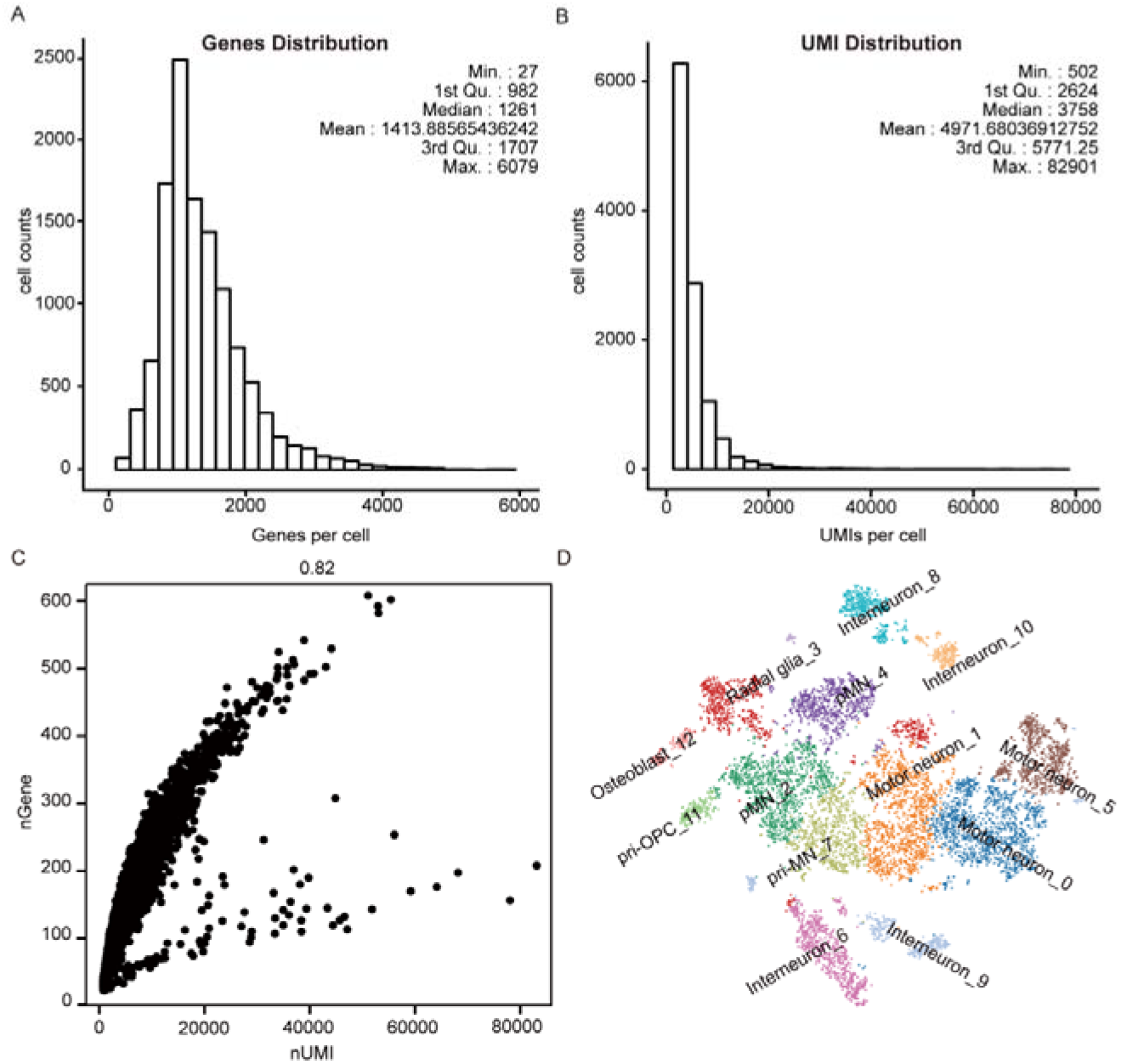
Technical performance and cell clustering of scRNA-seq. A-C) UMI counts or genes detected by scRNA-seq. D) tSNE visualization of cell clustering.

**Supplemental Figure 3:**
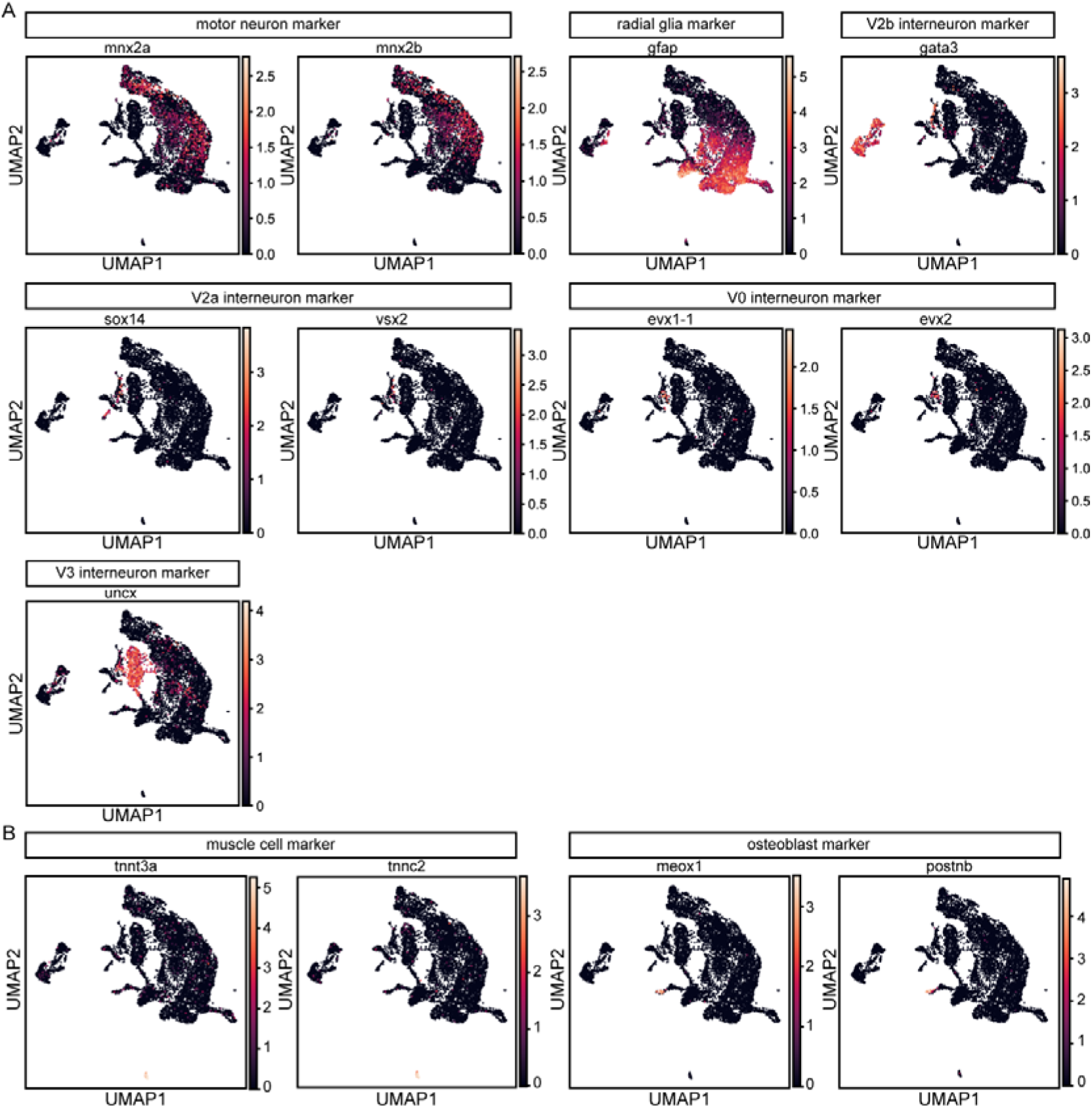
Identification of cell populations by known markers. UMAP visualization of known gene markers in interneurons, OPC lineages, motor neurons (A), muscle cells and osteoblasts (B).

**Supplemental Figure 4:**
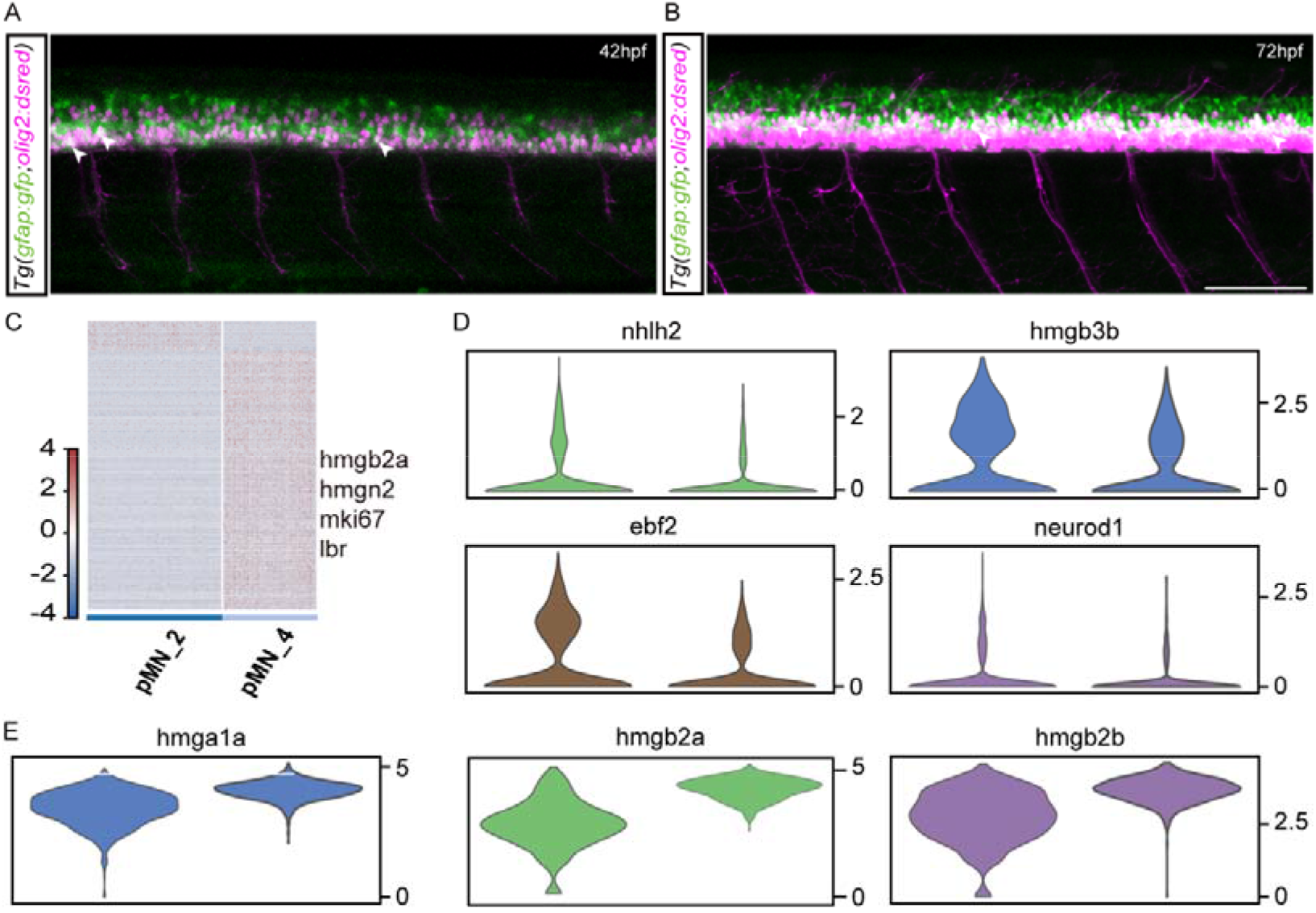
Enriched genes for pMN progenitor pMN_2 and pMN_4. A-B) pMN progenitor (olig2+) cells colabeled with GFAP. Confocal images of whole-mount *Tg(olig2:dsRed;GFAP:GFP)* embryos at 42 hpf (A) and 72 hpf (B), lateral view. C) Heatmaps of DEGs between pMN_2 and pMN_4. D-E) Violin plots of enriched genes in pMN_2 or pMN_4.

**Supplemental Figure 5:**
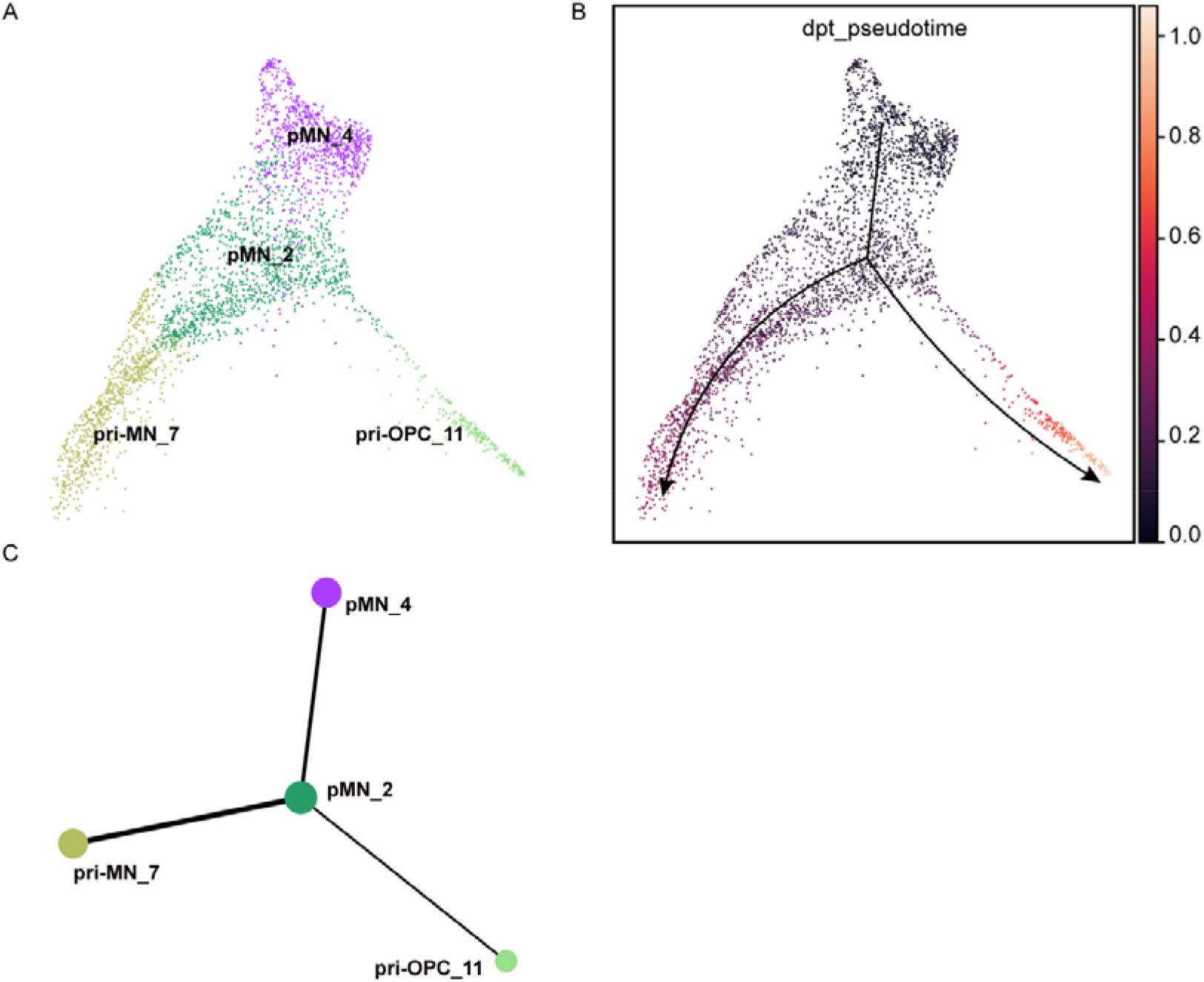
Pseudotemporal analysis of pMN progenitor or precursor subpopulation by PAGA.

**Supplemental Figure 6:**
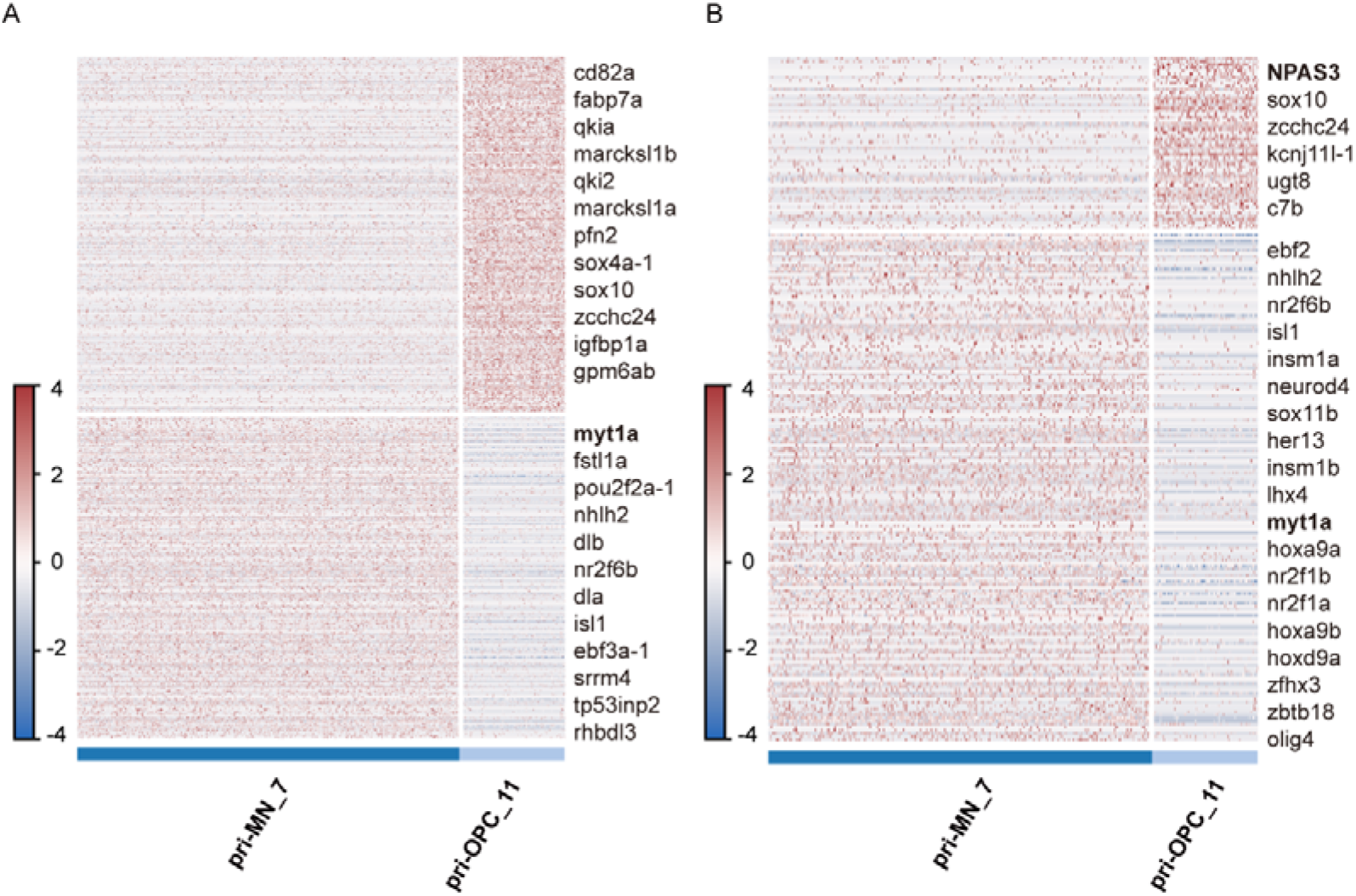
Heatmaps of differentially expressed genes or transcription factors between pri-MNs and pri-OPCs. Genes verified (*myt1* and *npas3*) are marked in bold.

**Supplemental Figure 7:**
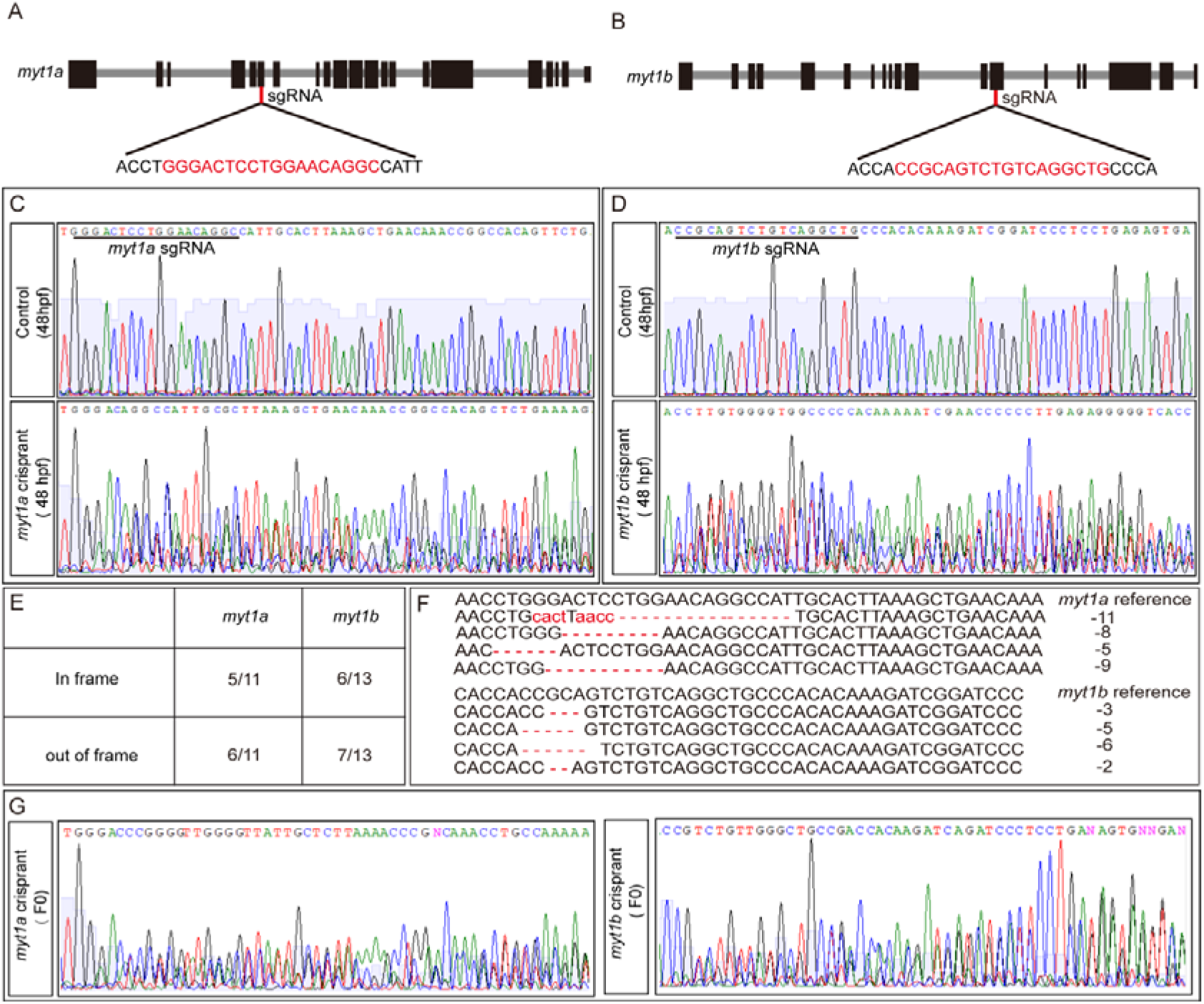
*myt1* including *myt1a* and *myt1b* mutagenesis generated by CRISPR. A-B) Schematic of *myt1a* and *myt1b* sgRNAs used for CRISPR. C-D) Representative Sanger sequencing results of *myt1* or *myt1b* somatic mutagenesis at 48 hpf. E-F) Gene editing types from individual *myt1* or *myt1b* PCR amplicons. G) Representative Sanger sequencing results of *myt1* or *myt1b* somatic mutagenesis in adult F0.

**Supplemental Figure 8:**
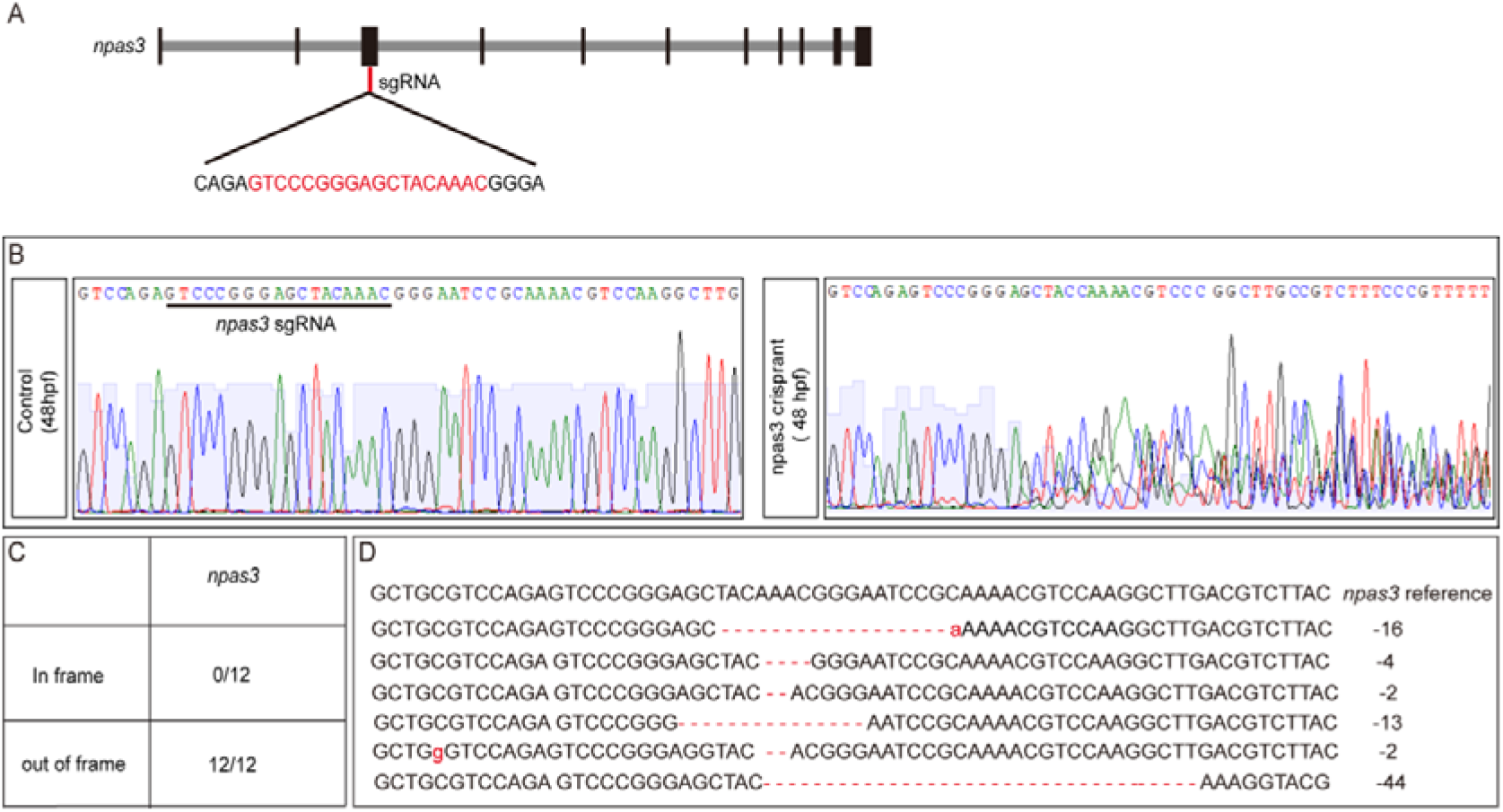
*npas3* mutagenesis generated by CRISPR. A) Schematic of the *npas3* sgRNA used for CRISPR. B) Representative Sanger sequencing results of *npas3* somatic mutagenesis in F0. C-D) Gene editing types from individual *npas3* PCR amplicons.

**Supplemental Figure 9:**
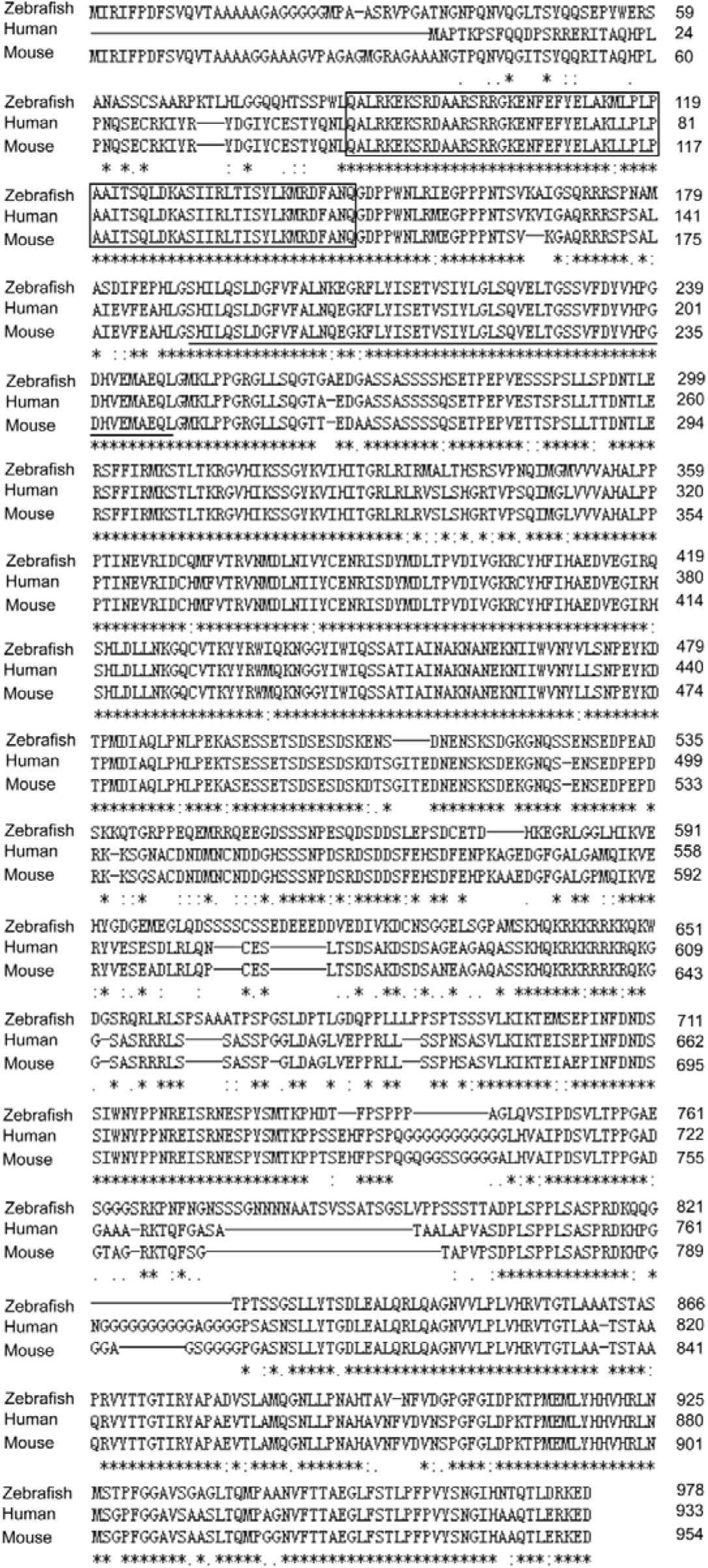
Amino acid alignment of NPAS3 in mouse, human, and zebrafish.

**Supplemental Figure 10:**
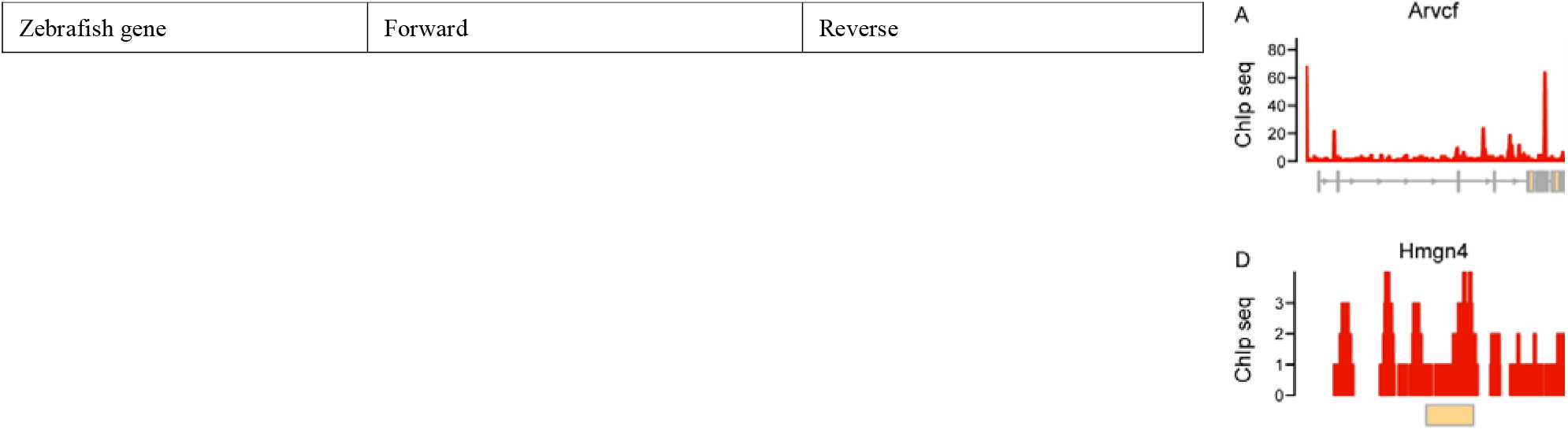
The binding profiles of NPAS3 and other “Schizophrenia Genes”.

**Supplemental Table 1:**
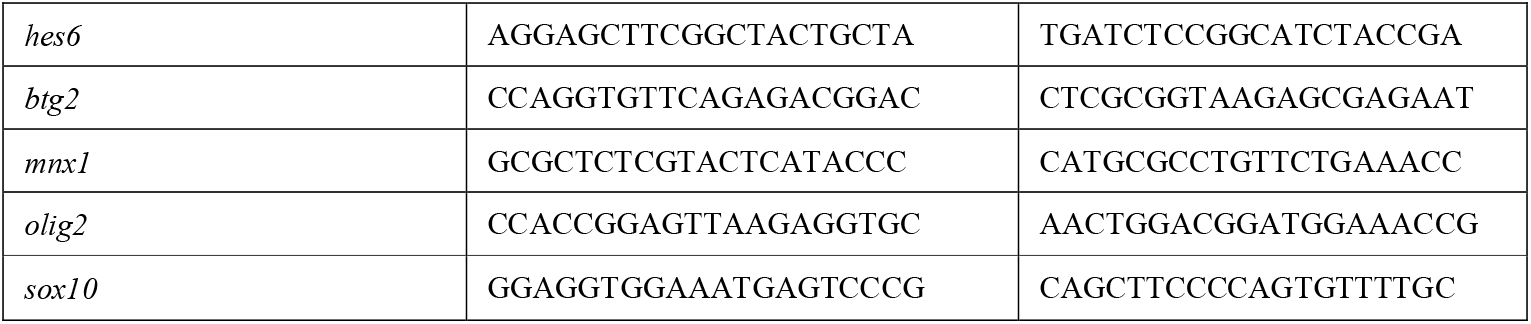
Primers for fluorescent *in situ* hybridization.

**Supplemental Table 2:** “Schizophrenia Genes” selected for this article. These genes found were associated with schizophrenia at least from two independent studies. 19 genes in bold indicate those detected in the dataset GSE70872. Red colors represent the 14 genes bound by the NPAS3 transcripton factor in the promoter regions or enhancer regions.

## References

Andrzejczuk, L. A., S. Banerjee, S. J. England, C. Voufo, K. Kamara, and K. E. Lewis. (2018). Tal1, Gata2a, and Gata3 Have Distinct Functions in the Development of V2b and Cerebrospinal Fluid-Contacting KA Spinal Neurons. Front Neurosci 12:170. doi: 10.3389/fnins.2018.00170.

Bayraktar, O. A., and C. Q. Doe. (2013). Combinatorial temporal patterning in progenitors expands neural diversity. Nature 498 7455:449–455. doi: 10.1038/nature12266.

Becht, E., L. McInnes, J. Healy, C. A. Dutertre, I. W. H. Kwok, L. G. Ng, F. Ginhoux, and E. W. Newell. (2018). Dimensionality reduction for visualizing single-cell data using UMAP. Nat Biotechnol. doi: 10.1038/nbt.4314.

Brunskill, E. W., L. A. Ehrman, M. T. Williams, J. Klanke, D. Hammer, T. L. Schaefer, R. Sah, G. W. Dorn, 2nd, S. S. Potter, and C. V. Vorhees. (2005). Abnormal neurodevelopment, neurosignaling and behaviour in Npas3-deficient mice. Eur J Neurosci 22 6:1265–1276. doi: 10.1111/j.1460-9568.2005.04291.x.

Carney, T. J., K. A. Dutton, E. Greenhill, M. Delfino-Machin, P. Dufourcq, P. Blader, and R. N. Kelsh. (2006). A direct role for Sox10 in specification of neural crest-derived sensory neurons. Development 133 23:4619–4630. doi: 10.1242/dev.02668.

Cleary, M. D., and C. Q. Doe. (2006). Regulation of neuroblast competence: multiple temporal identity factors specify distinct neuronal fates within a single early competence window. Genes Dev 20 4:429–434. doi: 10.1101/gad.1382206.

Clovis, Y. M., S. Y. Seo, J. S. Kwon, J. C. Rhee, S. Yeo, J. W. Lee, S. Lee, and S. K. Lee. (2016). Chx10 Consolidates V2a Interneuron Identity through Two Distinct Gene Repression Modes. Cell Rep 16 6:1642–1652. doi: 10.1016/j.celrep.2016.06.100.

Ding, S., Y. Gu, Y. Cai, M. Cai, T. Yang, S. Bao, W. Shen, X. Ni, G. Chen, and L. Xing. (2020). Integrative systems and functional analyses reveal a role of dopaminergic signaling in myelin pathogenesis. J Transl Med 18 1:109. doi: 10.1186/s12967-020-02276-1.

Farrell, J. A., Y. Wang, S. J. Riesenfeld, K. Shekhar, A. Regev, and A. F. Schier. (2018). Single-cell reconstruction of developmental trajectories during zebrafish embryogenesis. Science 360:eaar3131. doi: 10.1126/science.aar3131. Epub 2018 Apr 26.

Filtz, E. A., A. Emery, H. Lu, C. L. Forster, C. Karasch, and T. C. Hallstrom. (2015). Rb1 and Pten Co-Deletion in Osteoblast Precursor Cells Causes Rapid Lipoma Formation in Mice. PLoS One 10 8:e0136729. doi: 10.1371/journal.pone.0136729.

Flanagan-Steet, H., M. A. Fox, D. Meyer, and J. R. Sanes. (2005). Neuromuscular synapses can form in vivo by incorporation of initially aneural postsynaptic specializations. Development 132 20:4471–4481. doi: 10.1242/dev.02044.

Floriddia, E. M., T. Lourenco, S. Zhang, D. van Bruggen, M. M. Hilscher, P. Kukanja, J. P. Goncalves Dos Santos, M. Altinkok, C. Yokota, E. Llorens-Bobadilla, S. B. Mulinyawe, M. Graos, L. O. Sun, J. Frisen, M. Nilsson, and G. Castelo-Branco. (2020). Distinct oligodendrocyte populations have spatial preference and different responses to spinal cord injury. Nat Commun 11 1:5860. doi: 10.1038/s41467-020-19453-x.

Gejman, P. V., A. R. Sanders, and J. Duan. (2010). The role of genetics in the etiology of schizophrenia. Psychiatr Clin North Am 33 1:35–66. doi: 10.1016/j.psc.2009.12.003.

Gong, J., S. Hu, Z. Huang, Y. Hu, X. Wang, J. Zhao, P. Qian, C. Wang, J. Sheng, X. Lu, G. Wei, and D. Liu. (2020). The Requirement of Sox2 for the Spinal Cord Motor Neuron Development of Zebrafish. Front Mol Neurosci 13:34. doi: 10.3389/fnmol.2020.00034.

Guo, Z., L. Zhang, Z. Wu, Y. Chen, F. Wang, and G. Chen. (2014). In vivo direct reprogramming of reactive glial cells into functional neurons after brain injury and in an Alzheimer’s disease model. Cell Stem Cell 14 2:188–202. doi: 10.1016/j.stem.2013.12.001.

Hahne, F., and R. Ivanek. (2016). Visualizing Genomic Data Using Gviz and Bioconductor. Methods Mol Biol 1418:335–351. doi: 10.1007/978-1-4939-3578-9_16.

Henriksen, M. G., J. Nordgaard, and L. B. Jansson. (2017). Genetics of Schizophrenia: Overview of Methods, Findings and Limitations. Front Hum Neurosci 11:322. doi: 10.3389/fnhum.2017.00322.

Hu, J., J. Xu, L. Pang, H. Zhao, F. Li, Y. Deng, L. Liu, Y. Lan, X. Zhang, T. Zhao, C. Xu, C. Xu, Y. Xiao, and X. Li. (2016). Systematically characterizing dysfunctional long intergenic non-coding RNAs in multiple brain regions of major psychosis. Oncotarget 7:71087–71098. doi: 10.18632/oncotarget.12122.

Isshiki, T., B. Pearson, S. Holbrook, and C. Q. Doe. (2001). Drosophila neuroblasts sequentially express transcription factors which specify the temporal identity of their neuronal progeny. Cell 106:511–521. doi: 10.1016/s0092-8674(01)00465-2.

Juarez-Morales, J. L., C. J. Schulte, S. A. Pezoa, G. K. Vallejo, W. C. Hilinski, S. J. England, S. de Jager, and K. E. Lewis. (2016). Evx1 and Evx2 specify excitatory neurotransmitter fates and suppress inhibitory fates through a Pax2-independent mechanism. Neural Dev 11:5. doi: 10.1186/s13064-016-0059-9.

Kamnasaran, D., W. J. Muir, M. A. Ferguson-Smith, and D. W. Cox. (2003). Disruption of the neuronal PAS3 gene in a family affected with schizophrenia. J Med Genet 40 5:325–332. doi: 10.1136/jmg.40.5.325.

Kao, C. F., H. H. Yu, Y. He, J. C. Kao, and T. Lee. (2012). Hierarchical deployment of factors regulating temporal fate in a diverse neuronal lineage of the Drosophila central brain. Neuron 73 4:677–684. doi: 10.1016/j.neuron.2011.12.018.

Lange, C., F. Rost, A. Machate, S. Reinhardt, M. Lesche, A. Weber, V. Kuscha, A. Dahl, S. Rulands, and M. Brand. (2020). Single cell sequencing of radial glia progeny reveals the diversity of newborn neurons in the adult zebrafish brain. Development 147 1. doi: 10.1242/dev.185595.

Leber, S. M., S. M. Breedlove, and J. R. Sanes. (1990). Lineage, arrangement, and death of clonally related motoneurons in chick spinal cord. J Neurosci 10 7:2451–2462. doi: 10.1523/JNEUROSCI.10-07-02451.1990.

Leber, S. M., and J. R. Sanes. (1995). Migratory paths of neurons and glia in the embryonic chick spinal cord. J Neurosci 15:1236–1248. doi: 10.1523/JNEUROSCI.15-02-01236.1995.

Liang, X. G., C. Tan, C. K. Wang, R. R. Tao, Y. J. Huang, K. F. Ma, K. Fukunaga, M. Z. Huang, and F. Han. (2018). Myt1l induced direct reprogramming of pericytes into cholinergic neurons. CNS Neurosci Ther 24 9:801–809. doi: 10.1111/cns.12821.

Luoma, L. M., and F. B. Berry. (2018). Molecular analysis of NPAS3 functional domains and variants. BMC Mol Biol 19 1:14. doi: 10.1186/s12867-018-0117-4.

Mathews, E. S., and B. Appel. (2016). Oligodendrocyte differentiation. Methods Cell Biol 134:69–96. doi: 10.1016/bs.mcb.2015.12.004.

Metzis, V., S. Steinhauser, E. Pakanavicius, M. Gouti, D. Stamataki, K. Ivanovitch, T. Watson, T. Rayon, S. N. Mousavy Gharavy, R. Lovell-Badge, N. M. Luscombe, and J. Briscoe. (2018). Nervous System Regionalization Entails Axial Allocation before Neural Differentiation. Cell 175 4:1105–1118 e1117. doi: 10.1016/j.cell.2018.09.040.

Mizuguchi, R., M. Sugimori, H. Takebayashi, H. Kosako, M. Nagao, S. Yoshida, Y. Nabeshima, K. Shimamura, and M. Nakafuku. (2001). Combinatorial roles of olig2 and neurogenin2 in the coordinated induction of pan-neuronal and subtype-specific properties of motoneurons. Neuron 31 5:757–771. doi: 10.1016/s0896-6273(01)00413-5.

Moen, M. J., H. H. Adams, J. H. Brandsma, D. H. Dekkers, U. Akinci, S. Karkampouna, M. Quevedo, C. E. Kockx, Z. Ozgur, I. W. F. van, J. Demmers, and R. A. Poot. (2017). An interaction network of mental disorder proteins in neural stem cells. Transl Psychiatry 7 4:e1082. doi: 10.1038/tp.2017.52.

Nielsen, J. A., J. A. Berndt, L. D. Hudson, and R. C. Armstrong. (2004). Myelin transcription factor 1 (Myt1) modulates the proliferation and differentiation of oligodendrocyte lineage cells. Mol Cell Neurosci 25 1:111–123. doi: 10.1016/j.mcn.2003.10.001.

Ohnmacht, J., Y. Yang, G. W. Maurer, A. Barreiro-Iglesias, T. M. Tsarouchas, D. Wehner, D. Sieger, C. G. Becker, and T. Becker. (2016). Spinal motor neurons are regenerated after mechanical lesion and genetic ablation in larval zebrafish. Development 143 9:1464–1474. doi: 10.1242/dev.129155.

Park, H. C., J. Shin, and B. Appel. (2004). Spatial and temporal regulation of ventral spinal cord precursor specification by Hedgehog signaling. Development 131 23:5959–5969. doi: 10.1242/dev.01456.

Pickard, B. S., M. P. Malloy, D. J. Porteous, D. H. Blackwood, and W. J. Muir. (2005). Disruption of a brain transcription factor, NPAS3, is associated with schizophrenia and learning disability. Am J Med Genet B Neuropsychiatr Genet 136B 1:26–32. doi: 10.1002/ajmg.b.30204.

Pieper, A. A., X. Wu, T. W. Han, S. J. Estill, Q. Dang, L. C. Wu, S. Reece-Fincanon, C. A. Dudley, J. A. Richardson, D. J. Brat, and S. L. McKnight. (2005). The neuronal PAS domain protein 3 transcription factor controls FGF-mediated adult hippocampal neurogenesis in mice. Proc Natl Acad Sci U S A 102:14052–14057. doi: 10.1073/pnas.0506713102.

Qian, H., X. Kang, J. Hu, D. Zhang, Z. Liang, F. Meng, X. Zhang, Y. Xue, R. Maimon, S. F. Dowdy, N. K. Devaraj, Z. Zhou, W. C. Mobley, D. W. Cleveland, and X. D. Fu. (2020). Reversing a model of Parkinson’s disease with in situ converted nigral neurons. Nature 582 7813:550–556. doi: 10.1038/s41586-020-2388-4.

Qian, X., S. K. Goderie, Q. Shen, J. H. Stern, and S. Temple. (1998). Intrinsic programs of patterned cell lineages in isolated vertebrate CNS ventricular zone cells. Development 125 16:3143–3152.

Qian, X., Q. Shen, S. K. Goderie, W. He, A. Capela, A. A. Davis, and S. Temple. (2000). Timing of CNS cell generation a programmed sequence of neuron and glial cell production from isolated murine cortical stem cells. Neuron 28 1:69–80. doi: 10.1016/s0896-6273(00)00086-6.

Ravanelli, A. M., and B. Appel. (2015). Motor neurons and oligodendrocytes arise from distinct cell lineages by progenitor recruitment. Genes Dev 29 23:2504–2515. doi: 10.1101/gad.271312.115.

Reimer, M. M., A. Norris, J. Ohnmacht, R. Patani, Z. Zhong, T. B. Dias, V. Kuscha, A. L. Scott, Y. C. Chen, S. Rozov, S. L. Frazer, C. Wyatt, S. Higashijima, E. E. Patton, P. Panula, S. Chandran, T. Becker, and C. G. Becker. (2013). Dopamine from the brain promotes spinal motor neuron generation during development and adult regeneration. Dev Cell 25 5:478–491. doi: 10.1016/j.devcel.2013.04.012.

Richardson, W. D., H. K. Smith, T. Sun, N. P. Pringle, A. Hall, and R. Woodruff. (2000). Oligodendrocyte lineage and the motor neuron connection. Glia 29 2:136–142. doi: 10.1002/(sici)1098-1136(20000115)29:2<136::aid-glia6>3.0.co;2-g.

Schmid, A., A. Chiba, and C. Q. Doe. (1999). Clonal analysis of Drosophilaembryonic neuroblasts: neural cell types, axon projections and muscle targets. Development 126:4653–4689.

Scott, K., R. O’Rourke, A. Gillen, and B. Appel. (2020). Prdm8 regulates pMN progenitor specification for motor neuron and oligodendrocyte fates by modulating the Shh signaling response. Development 147 16. doi: 10.1242/dev.191023.

Seredick, S. D., L. V. Ryswyk, S. A. Hutchinson, and J. S. Eisen. (2012). Zebrafish Mnx proteins specify one motoneuron subtype and suppress acquisition of interneuron characteristics. Neural Dev 7:35. doi: 10.1186/1749-8104-7-35.

Shin, J., H.-C. Park, J. M. Topczewska, D. J. Mawdsley, and B. Appel. (2003). Neural cell fate analysis in zebrafish using olig2 BAC transgenics. Methods Cell Sci 25:7–14. doi: 10.1023/B:MICS.0000006847.09037.3a.

Takada, N., and B. Appel. (2010). Identification of genes expressed by zebrafish oligodendrocytes using a differential microarray screen. Dev Dyn 239 7:2041–2047. doi: 10.1002/dvdy.22338.

Takahashi, N., T. Sakurai, K. L. Davis, and J. D. Buxbaum. (2011). Linking oligodendrocyte and myelin dysfunction to neurocircuitry abnormalities in schizophrenia. Prog Neurobiol 93 1:13–24. doi: 10.1016/j.pneurobio.2010.09.004.

Terzi, Y. K., S. Oğuzkan-Balci, B. Anlar, E. Erdoğan-Bakar, and S. Ayter. (2011). Learning disability and oligodendrocyte myelin glycoprotein (OMGP) gene in neurofibromatosis type 1. Turk J Pediatr 53 1:75–78.

Thyme, S. B., L. M. Pieper, E. H. Li, S. Pandey, Y. Wang, N. S. Morris, C. Sha, J. W. Choi, K. J. Herrera, E. R. Soucy, S. Zimmerman, O. Randlett, J. Greenwood, S. A. McCarroll, and A. F. Schier. (2019). Phenotypic Landscape of Schizophrenia-Associated Genes Defines Candidates and Their Shared Functions. Cell 177 2:478–491 e420. doi: 10.1016/j.cell.2019.01.048.

Trapnell, C., D. Cacchiarelli, J. Grimsby, P. Pokharel, S. Li, M. Morse, N. J. Lennon, K. J. Livak, T. S. Mikkelsen, and J. L. Rinn. (2014). The dynamics and regulators of cell fate decisions are revealed by pseudotemporal ordering of single cells. Nat Biotechnol 32 4:381–386. doi: 10.1038/nbt.2859.

Vasconcelos, F. F., A. Sessa, C. Laranjeira, A. Raposo, V. Teixeira, D. W. Hagey, D. M. Tomaz, J. Muhr, V. Broccoli, and D. S. Castro. (2016). MyT1 Counteracts the Neural Progenitor Program to Promote Vertebrate Neurogenesis. Cell Rep 17 2:469–483. doi: 10.1016/j.celrep.2016.09.024.

Vierbuchen, T., A. Ostermeier, Z. P. Pang, Y. Kokubu, T. C. Sudhof, and M. Wernig. (2010). Direct conversion of fibroblasts to functional neurons by defined factors. Nature 463 7284:1035–1041. doi: 10.1038/nature08797.

Wagner, D. E., C. Weinreb, Z. M. Collins, J. A. Briggs, S. G. Megason, and A. M. Klein. (2018). Single-cell mapping of gene expression landscapes and lineage in the zebrafish embryo. Science 360:981–987. doi: 10.1126/science.aar4362. Epub 2018 Apr 26.

Weinberger, M., F. C. Simoes, R. Patient, T. Sauka-Spengler, and P. R. Riley. (2020). Functional Heterogeneity within the Developing Zebrafish Epicardium. Dev Cell 52 5:574–590 e576. doi: 10.1016/j.devcel.2020.01.023.

Weng, Q., J. Wang, J. Wang, D. He, Z. Cheng, F. Zhang, R. Verma, L. Xu, X. Dong, Y. Liao, X. He, A. Potter, L. Zhang, C. Zhao, M. Xin, Q. Zhou, B. J. Aronow, P. J. Blackshear, J. N. Rich, Q. He, W. Zhou, M. L. Suva, R. R. Waclaw, S. S. Potter, G. Yu, and Q. R. Lu. (2019). Single-Cell Transcriptomics Uncovers Glial Progenitor Diversity and Cell Fate Determinants during Development and Gliomagenesis. Cell Stem Cell 24 5:707–723 e708. doi: 10.1016/j.stem.2019.03.006.

Wolf, F. A., P. Angerer, and F. J. Theis. (2018). SCANPY: large-scale single-cell gene expression data analysis. Genome Biol 19 1:15. doi: 10.1186/s13059-017-1382-0.

Xie, Y., D. Kaufmann, M. J. Moulton, S. Panahi, J. A. Gaynes, H. N. Watters, D. Zhou, H. H. Xue, C. M. Fung, E. M. Levine, A. Letsou, K. C. Brennan, and R. I. Dorsky. (2017). Lef1-dependent hypothalamic neurogenesis inhibits anxiety. PLoS Biol 15 8:e2002257. doi: 10.1371/journal.pbio.2002257.

Xing, L., J. H. Son, T. J. Stevenson, C. Lillesaar, L. Bally-Cuif, T. Dahl, and J. L. Bonkowsky. (2015). A Serotonin Circuit Acts as an Environmental Sensor to Mediate Midline Axon Crossing through EphrinB2. J Neurosci 35 44:14794–14808. doi: 10.1523/JNEUROSCI.1295-15.2015.

Yang, J., Y. H. Shih, and X. Xu. (2014). Understanding cardiac sarcomere assembly with zebrafish genetics. Anat Rec (Hoboken) 297 9:1681–1693. doi: 10.1002/ar.22975.

Yang, T., L. Xing, W. Yu, Y. cai, S. Cui, and G. Chen. (2020). Astrocytic reprogramming combined with rehabilitation strategy improves recovery from spinal cord injury. FASEB J 34 11:15504–15515. doi: 10.1096/fj.202001657RR. Epub 2020 Sep 25.

Zheng, G. X., J. M. Terry, P. Belgrader, P. Ryvkin, Z. W. Bent, R. Wilson, S. B. Ziraldo, T. D. Wheeler, G. P. McDermott, J. Zhu, M. T. Gregory, J. Shuga, L. Montesclaros, J. G. Underwood, D. A. Masquelier, S. Y. Nishimura, M. Schnall-Levin, P. W. Wyatt, C. M. Hindson, R. Bharadwaj, A. Wong, K. D. Ness, L. W. Beppu, H. J. Deeg, C. McFarland, K. R. Loeb, W. J. Valente, N. G. Ericson, E. A. Stevens, J. P. Radich, T. S. Mikkelsen, B. J. Hindson, and J. H. Bielas. (2017). Massively parallel digital transcriptional profiling of single cells. Nat Commun 8:14049. doi: 10.1038/ncomms14049.

Zhou, Q., G. Choi, and D. J. Anderson. (2001). The bHLH transcription factor Olig2 promotes oligodendrocyte differentiation in collaboration with Nkx2.2. Neuron 31 5:791–807. doi: 10.1016/s0896-6273(01)00414-7.

